# Rapid and in-depth coverage of the (phospho-)proteome with deep libraries and optimal window design for dia-PASEF

**DOI:** 10.1101/2022.05.31.494163

**Authors:** Patricia Skowronek, Marvin Thielert, Eugenia Voytik, Maria C. Tanzer, Fynn M. Hansen, Sander Willems, Özge Karayel, Andreas-David Brunner, Florian Meier, Matthias Mann

## Abstract

Data-independent acquisition (DIA) methods have become increasingly attractive in mass spectrometry (MS)-based proteomics, because they enable high data completeness and a wide dynamic range. Recently, we combined DIA with parallel accumulation – serial fragmentation (dia-PASEF) on a Bruker trapped ion mobility separated (TIMS) quadrupole time-of-flight (TOF) mass spectrometer. This requires alignment of the ion mobility separation with the downstream mass selective quadrupole, leading to a more complex scheme for dia-PASEF window placement compared to DIA. To achieve high data completeness and deep proteome coverage, here we employ variable isolation windows that are placed optimally depending on precursor density in the *m/z* and ion mobility plane. This <u>A</u>utomatic Isolation <u>D</u>esign procedure is implemented in the freely available py_diAID package. In combination with in-depth project-specific proteomics libraries and the Evosep LC system, we reproducibly identified over 7,700 proteins in a human cancer cell line in 44 minutes with quadruplicate single-shot injections at high sensitivity. Even at a throughput of 100 samples per day (11 minutes LC gradients), we consistently quantified more than 6,000 proteins in mammalian cell lysates by injecting four replicates. We found that optimal dia-PASEF window placement facilitates in-depth phosphoproteomics with very high sensitivity, quantifying more than 35,000 phosphosites in a human cancer cell line stimulated with an epidermal growth factor (EGF) in triplicate 21 minutes runs. This covers a substantial part of the regulated phosphoproteome with high sensitivity, opening up for extensive systems-biological studies.

## INTRODUCTION

MS-based proteomics has become a powerful tool to study proteomes in a systematic and unbiased manner (1). In recent years, this development has been accelerated by data-independent acquisition (DIA) (2), where predefined isolation windows cycle through the *m/z*-range of interest, and regularly subject the covered peptide precursors to fragmentation (3– 6). Although the concept of DIA was established more than a decade ago (4, 7), only the most recent DIA implementations and hardware advancements in MS and data analysis are at par or even exceeding data dependent acquisition (DDA) with regards to sensitivity, reproducibility, and dynamic range coverage (2, 6, 8) and surpass targeted approaches in throughput and ease-of-use (9, 10). This holds also true for studying post-translational modifications (11–13).

DIA has recently shown promise in combination with trapped ion mobility spectrometry (TIMS) mass spectrometers, as demonstrated with single-cell analysis (14, 15). The TIMS tunnel is a compact and high-performance implementation of ion mobility separation. It captures the peptides from the incoming ion beam discretizing the continuous LC elution. Within the TIMS tunnel, each ion reaches an equilibrium position based on the opposing forces of a gas flow and an electric field gradient. Decreasing the electric field gradient elutes the peptide ions as a function of their ion mobility (16–19). In the Bruker timsTOF instruments, the TIMS device is placed upstream of mass-selective quadrupole and high-resolution time-of-flight mass analyzer and is itself divided into two parts (20–22). The mobility separation can be synchronized with the quadrupole isolation, leading to high ion beam utilization, increased sensitivity and decreased spectral complexity due to the additional ion mobility dimension (6, 20, 23). This principle is termed PASEF for parallel accumulation-serial fragmentation (21, 24).

When combined with DIA (dia-PASEF), peptide precursors separate not only in the *m/z* but also in the ion mobility dimension, in contrast to standard DIA modes (2, 6). We have observed that dia-PASEF is particularly beneficial for acquiring a wide range of proteomics data while maintaining a high sequence coverage and very high sensitivity (6, 15). Furthermore, ions are detected by inherently fast TOF analysis allowing fast DIA cycle times, which is particularly advantageous for short LC gradients (6). The resulting, complex spectra can be efficiently analyzed by machine learning or deep learning-based algorithms such as DIA-NN (25, 26). Here, we set out to explore the potential of dia-PASEF to further increase coverage and quantitative accuracy on the fast and sensitive ion mobility-mass spectrometry platform. In dia-PASEF, two-dimensional precursor isolation schemes are defined in the *m/z*-ion mobility plane. We used a Bayesian optimization algorithm ensuring optimal placement of the acquisition scheme in both dimensions. Single-runs acquired with these optimal dia-PASEF methods were searched against in-depth project-specific libraries. Furthermore, we combined dia-PASEF with the Evosep One LC system, which features a pre-formed gradient particularly designed for high throughput by eliminating inter-run overhead (6, 27). Together, our optimized dia-PASEF workflow for high throughput proteomics quantified more than 7,000 proteins in only 21 minutes from quadruplicate injections of a tryptic HeLa digest. Motivated by these proteomic results, we also investigated py_diAID for phosphorylation analysis. On the Orbitrap MS platform, Olsen and co-workers recently demonstrated an efficient combination of fast chromatography runs with DIA, quantifying more than 13,000 phosphopeptides in very short (15 min) LC/MS runs from HeLa cells using the Spectronaut software (11). In a small scale study, Ishihama and co-workers showed that phosphopeptides analysis benefits from the additional ion mobility dimension in PASEF (28). For large-scale PTM studies, our optimized py_diAID acquisition schemes cover nearly all theoretical phosphopeptide precursors and quantified expected changes in the well-studied EGF-receptor signaling pathway with minimal time and sample consumption.

## EXPERIMENTAL PROCEDURES

### Experimental Design and Statistical Rationale

All experiments were done using HeLa cell lysate obtained from HeLa S3 cells (ATCC), routinely used for proteomics method development and benchmark experiments (supplemental Fig. S1). Altogether, the data set includes 322 raw data files (uploaded to PRIDE, see below). We used the same HeLa batch for generating libraries and single-run data of both proteome and phosphoproteome measurements. In brief, proteome measurements with different gradient lengths and the technical comparisons of the original and optimal dia-PASEF methods for phosphoproteomics were acquired in quadruplicates. Unless otherwise mentioned, 200 ng HeLa lysate were used for single-run proteome and 100 μg for the single-run phoshopeptide enrichment experiments. The libraries were acquired as described below. The experimental design and statistical rational are described in the respective figure legends. The EGF experiment was performed in biological triplicates to determine significantly different phosphosite levels between the EGF-treated and control samples. Technical quadruplicates were acquired to evaluate reproducibility and quantitative accuracy by calculating coefficient of variations (CVs) and mean of the replicate injections. Moreover, we alternated the MS run order to avoid potential carryover effects or any similar biases.

### Sample preparation

HeLa S3 cells (ATCC) were cultured in Dulbecco’s modified Eagle’s medium (Life Technologies Ltd., UK) containing 20 mM glutamine, 10% fetal bovine serum, and 1% penicillin-streptomycin. Sample preparation was essentially performed as previously described in the in-stage tip protocol (29). In brief, the cells were washed with PBS and lysed. Protein reduction and alkylation and digestion with trypsin (Sigma-Aldrich) and LysC (WAKO) (1:100, enzyme/protein, w/w) were performed in one step. Resulting peptides were dried and reconstituted in a solution A* (0.1% TFA/2% ACN). Peptide concentrations were measured optically at 280 nm (Nanodrop 2000; Thermo Scientific) and 200 ng peptides were loaded onto Evotips for LC-MS/MS analysis as described previously (15). The Evotips were washed with 0.1% FA/99.9% ACN, equilibrated with 0.1% FA, loaded with the sample dissolved in 0.1% FA, and washed with 0.1% FA.

For phosphoproteomics, HeLa cells at a plate confluence of 80% were treated for 10 min with 100 ng/mL animal-free recombinant human EGF (PeproTech) or Gibco™ distilled water (Thermo Fisher Scientific) and washed three times with ice-cold TBS before lysis in 2% SDC in 100 mM Tris-HCl (pH 8.5) at 95°C. Protein concentrations were determined using the BCA assay and samples were then reduced and alkylated with 10 mM TCEP and 40 mM CAA, respectively. Altogether, 25 mg protein material of sample was used for the library generation, 8 mg for EGF treated experiments including method benchmarking and 4 mg for untreated experiments. The sample was digested with trypsin (Sigma-Aldrich) and LysC (WAKO) (1:100, enzyme/protein, w/w) overnight and subsequently desalted using Sepax Extraction columns (Generik DBX). Each cartridge was prepared with 100% MeOH and 99% MeOH/1% TFA. After equilibration with 0.2% TFA, the samples were loaded with a protein concentration of 1 mg/mL, washed with 99% IPA/1% TFA, 0.2% TFA/5% ACN, and 0.2% TFA solutions. The peptides were eluted with 5% NH_4_OH/80% ACN. Lyophilized peptides were reconstituted in equilibration solution (1% TFA/80% ACN) and 100 μg peptide material per sample/AssayMAP cartridge, each containing 5 μL Fe(III)-NTA, was enriched for phosphopeptide with the AssayMAP bravo robot (Agilent) (30). Phosphopeptides were dried in a SpeedVac for 20 min at 45°C and loaded onto Evotips as described above.

### High-pH reversed-phase fractionation for library generation

To generate proteome libraries, 10 μg and 60 μg peptides were separated with high pH reverse-phase chromatography into 24 and 48 fractions, respectively, on a 30 cm C_18_ column with an inner diameter of 250 μm at a flow rate of 2 μL/min using the spider sample fractionator (31). The gradient consisted of the binary buffer system (PreOmics GmbH). The buffer B concentration of 3% was increased to 30% in 45 min, 40% in 12 min and 60% in 5 min, and 95% in 10 min. After washing at 95% for 10 min, buffer B concentration was re-equilibrated to 3% in 10 min. The exit valve concatenated the eluted peptides automatically by switching after a defined collection time (80s for 24 and 60s for 48 fractions). The fractions were dried in a SpeedVac and reconstituted in solution A*. A quarter of each fraction was loaded onto Evotips for LC-MS/MS analysis. Below we will refer to ‘the reference proteome library’ that represents a 24 high pH fractions and dda-PASEF spectral library of a tryptic HeLa digest acquired with a 21 min Evosep gradient.

To generate a phosphoproteome library, peptides obtained from the EGF stimulated cells were separated using an UFLC system (Shimadzu). 6 mg peptide material was fractionated with a binary buffer system: A (2.5 mM ABC) and B (2.5 mM ABC/80% ACN). The peptides were loaded onto a reversed-phase column (ZORBAX 300Extend-C_18_, Agilent) and separated at a 1 mL/min flow rate at 40°C. The buffer B concentration of 2.5% was increased to 38% in

82.5 min, 75% in 2 min, and 100% in 8 min. It stayed at 100% for 2 min and was reduced to 2.5% in 2 min. In total, 95 fractions were collected and fractions with low peptide yield, as determined using Nanodrop, were pooled (supplemental table 1) and dried in a SpeedVac. Next, 76 fractions were enriched for phosphopeptide, which were subsequently loaded onto Evotips.

### LC-MS/MS analysis

The Evosep One liquid chromatography system coupled with a timsTOF Pro mass spectrometer (Bruker) was used to measure all samples. The 60 and 100 SPD (samples per day) methods required an 8 cm × 150 μm reverse-phase column packed with 1.5 μm C_18_-beads (Pepsep) and the 30 SPD method a 15 cm × 75 μm column with 1.9 μm C_18_-beads (Pepsep) at 40°C. The analytical columns were connected with a fused silica ID emitter (10 μm ID, Bruker Daltonics) inside a nano-electrospray ion source (Captive spray source, Bruker). The mobile phases comprised 0.1% FA as solution A and 0.1% FA/80% ACN as solution B.

The library samples were acquired in dda-PASEF mode with four PASEF/MSMS scans at a throughput of 60 and 100 SPDs and 10 PASEF/MSMS scans at 30 SPD per topN acquisition cycle. Singly charged precursors were filtered out by their position in the *m/z*-ion mobility plane, and only precursor signals over an intensity threshold of 2,500 arbitrary units (a.u.) were picked for fragmentation. While precursors over the target value of 20,000 a.u. were dynamically excluded for 0.4 min, ones below 700 Da were isolated with a 2 Th window and ones above with 3 Th. All spectra were acquired within an *m/z*-range of 100 to 1700 and an ion mobility range from 1.51 to 0.6 Vs cm^-2^.

We described the original dia-PASEF method in Meier et al. (6). The dia-PASEF methods optimized here with py_diAID cover an *m/z*-range from 300 to 1200 for proteome and from 400 to 1400 for phosphoproteome measurements. Each method includes two ion mobility windows per dia-PASEF scan with variable isolation window widths adjusted to the precursor densities. Eight, 12 and 25 dia-PASEF scans were deployed at a throughput of 100 (cycle time: 0.96 s), 60 (cycle time: 1.38s), and 30 SPDs (cycle time: 2.7s), respectively. We created dia-PASEF methods with equidistant window widths (supplemental Fig. S5) with the software “Compass DataAnalysis” (Bruker Daltonics). These acquisition schemes are plotted on top on a kernel density estimation of precursors from a reference library in supplemental Figure S2-4. The ion mobility range was set to 1.5 Vs cm^-2^ and 0.6 Vs cm^-2^. The accumulation and ramp times were specified as 100 ms for all experiments. As a result, each MS1 scan and each MS2/dia-PASEF scan last 100 ms plus additional transfer time, and a dia-PASEF method with 12 dia-PASEF scans has a cycle time of 1.38s. The collision energy was decreased as a function of the ion mobility from 59 eV at 1/K_0_ = 1.6 Vs cm^-2^ to 20 eV at 1/K_0_ = 0.6 Vs cm^-2^ and the ion mobility dimension was calibrated with three Agilent ESI Tuning Mix ions (*m/z*, 1/K_0_: 622.02,0.98 Vs cm^-2^, 922.01, 1.19 Vs cm^-2^, 1221.99, 1.38 Vs cm^-2^). For phosphoproteomics experiments, the collision energy was decreased from 60 eV at 1.5 Vs cm^-2^ to 54 eV at 1.17 Vs cm^-2^ to 25 eV at 0.85 Vs cm^-2^ and end at 20 eV at 0.6 Vs cm^-2^.

### Raw data analysis

We employed DIA-NN, MSFragger and Spectronaut for transforming raw data into precursor and fragment identifications based on 3D peak position (RT, *m/z* precursor, and ion mobility). In each case, all data was searched against the reviewed human proteome (Uniprot, Nov 2021, 20,360 entries without isoforms) with trypsin/LysC as digestion enzymes. Cysteine carbamidomethylation was set as fixed modification. Methionine oxidation, methionine excision at the N-terminus, and in the case of the phosphoproteome searches, phosphorylation (STY) was selected as variable modifications. A maximum of two missed cleavages and up to three variable modifications were allowed.

The project-specific libraries for DIA-NN analyses were generated with FragPipe (32) (FragPipe 16.2, MSFragger 3.4 (33–35), Philosopher 4.0.0 (36), Python 3.8, EasyPQP 0.1.25 (37)). The default settings were kept except that the precursor mass tolerance was set from -20 to 20 ppm and the fragment mass tolerance to 20 ppm. Additionally, Pyro-Glu or ammonia loss at the peptide N-terminus and water loss on N-terminal glutamic acid were selected as variable modification. The output tables were filtered for an 1% FDR using the Percolator (38, 39) and ProteinProphet (40) option in FragPipe (supplemental table 2).

DIA-NN 1.8 was used to analyze the single-shot experiments against the project-specific libraries generated with FragPipe (32). The default settings were kept except that we changed the charge state to 2 -4. The precursor’s *m/z* range was restricted from 300 to 1200 for proteome and 400 to 1400 for phosphoproteome analysis. The fragment *m/z* range was set from 100 to 1700, and the mass and MS1 accuracy to 15 ppm. ‘Match between run’ was enabled while ‘protein inference’ was disabled. We also enabled ‘robust LC (high precision)’ as the quantification strategy. The proteomics output tables were filtered for a maximum of 1% of q-value at both precursor and global protein levels. For phosphoproteomics, the post-translational modification q-value also had to be a maximum of 1%. The PG.MaxLFQ column integrated in the DIA-NN output tables reports normalized quantity employing the MaxLFQ principle (41) and was used for quantitative analysis on the protein level. For our phosphoproteomics analysis, we used the scoring of post-translational sites implemented in DIA-NN with ‘PTM.Site.Confidence’ indicating the localization probability (13).

Spectronaut (v16, Biognosys AG, Schlieren, Switzerland) (3) was used for comparative analysis and we used the same search settings as described above if not stated differently. The FDR cutoff was set to 1%. The precursor peptide and q-value cutoffs were 0.2 and 0.01, respectively. The protein q-value experiment and run wide cutoffs were 0.01 and 0.05, respectively. The dataset was analyzed with a sparse q-value and no imputation was performed. For phosphoproteomics experiments, the PTM localization cutoff was set to 0. The results were filtered for the best N fragments per peptide between 3 to 25.

Peptide collapse (v1.4.1), a plug-in tool for Perseus (42), collapsed peptide output tables from DIA-NN or Spectronaut to phosphosite tables using default settings and a localization cutoff of 0.75 (Class I sites) (11). The DIA-NN output table was reformatted by renaming all columns and entries calculating peptide positions to conform to the format required for the plug-in tool. For collapsing, Perseus took only phosphorylation into account. During collapsing phosphopeptide ions to phosphosites, each phosphosite corresponding to the same peptide obtains the same intensity, however imputation may lead to differences in fold changes. If the same phosphosite was identified on different peptides, which may also have modifications other than phosphorylation or different charge states, the intensities were summed up.

## Statistical Analysis

Visualization and statistical analyses were performed using the output tables of DIA-NN or Spectronaut with Python (3.8, Jupyter notebook) and the packages pandas (1.4.2) and pyfaidx (0.6.1) for data accession and py_diAID (0.0.16), AlphaMap (0.1.10), matplotlib (3.4.3), and seaborn (0.11.2) for visualization. The statistical analysis of the EGF experiment was performed in Perseus (1.6.2.2). Log_2_-transformed intensities were filtered for 100 % valid values in at least one condition. The missing values were replaced drawing from a normal distribution (width 0.3 and downshift 1.8). Next, we applied the two-sided Student’s t-test (S_0_=0.1, FDR = 0.05) to obtain the significantly changing phosphorylated peptides. A Fisher’s exact test was performed for GO term and KEGG pathway enrichment analysis (p-value<0.002).

## RESULTS

### Principle and limitations of the original dia-PASEF window design

In the timsTOF mass spectrometer (Bruker Daltonics), a dual TIMS tunnel releases the captured peptide ion species individually as a function of their mobility. In a PASEF MS/MS scan, a quadrupole transmits part of the ion beam where the precursor *m/z* values fall into a pre-defined isolation window (Fig. 1A). These precursors are subsequently fragmented by applying a particular collision energy. A downstream TOF analyzer acquires high-resolution mass spectra. In dia-PASEF, changing the quadrupole position is synchronized to the ion mobility elution, increasing the MS efficiency because the isolation window is placed on top of the precursor cloud (6). This movement happens in distinct steps and thereby divides one PASEF scan into multiple ion mobility windows. The quadrupole isolation window is first placed at high *m/z* for a certain amount of time, after which it jumps to a position in the lower *m/z* range. This transition point corresponds to a particular ion mobility value for each dia-PASEF scan. In each subsequent dia-PASEF scan, the starting *m/z* window is offset to lower values (Fig. 1B, C). Together, these isolation windows cover a large proportion of the *m/z* and the ion mobility dimensions, constituting a two-dimensional acquisition scheme (Fig. 1B).

**Figure 1:**
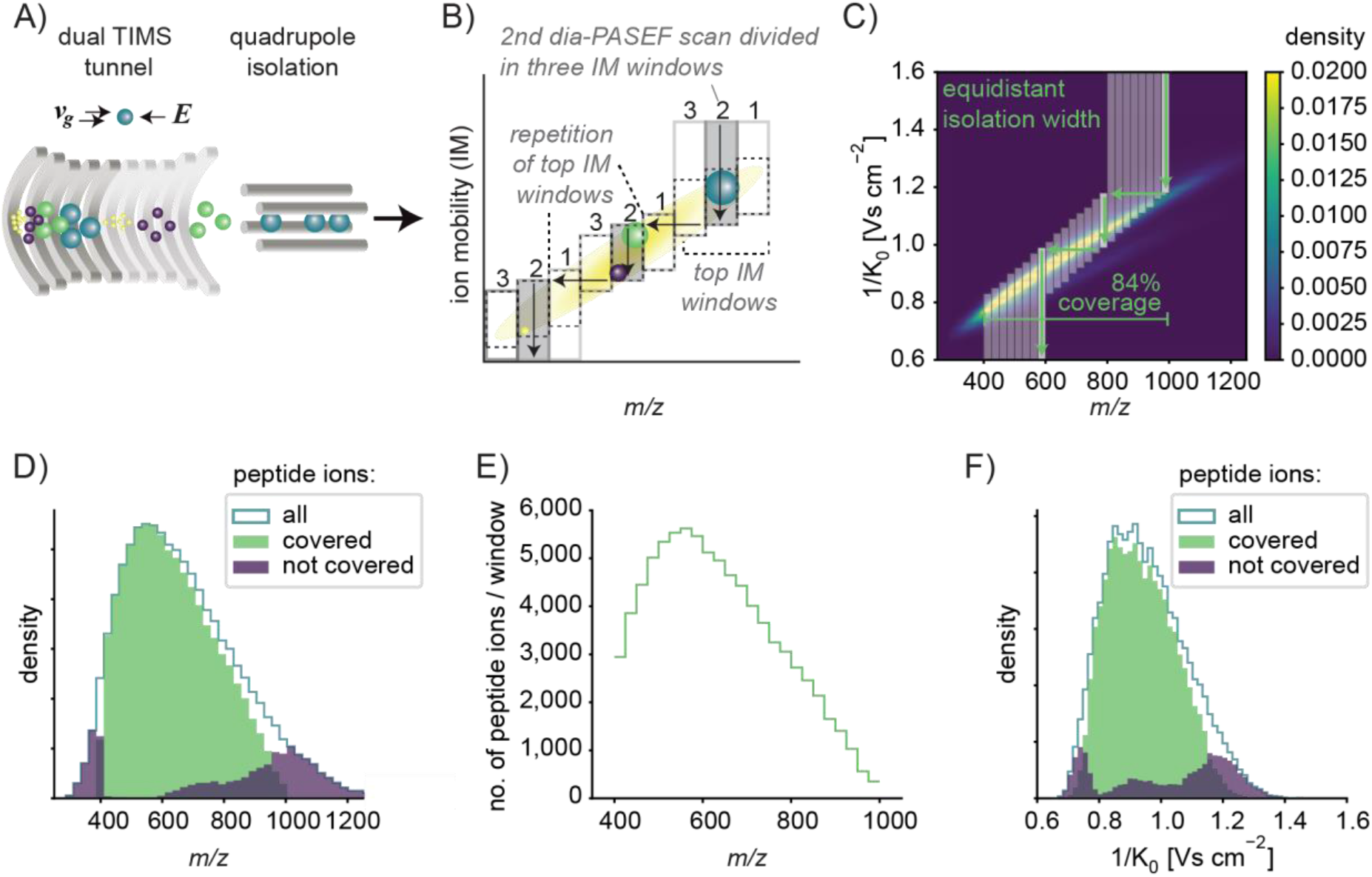
Principle of dia-PASEF on a timsTOF with equidistant two-dimensional isolation windows. A) Schematic of a TIMS tunnel followed by quadrupole isolation B) dia-PASEF acquisition scheme depicting three dia-PASEF scans divided into three ion mobility (IM) windows. Vertical arrows indicate the elution of the ions with decreasing electrical field and horizontal arrows indicate the movement of the quadrupole. The pattern of the top ion mobility windows is repeated and the top and bottom ion mobility windows are extended to the upper and lower ion mobility range, respectively. C) Original dia-PASEF acquisition scheme (6) plotted on a kernel density distribution of all precursors. One dia-PASEF scan is divided into three ion mobility windows by three distinct movements of quadrupole isolation. This scheme comprises eight dia-PASEF scans with equidistant isolation width covering in total 84% of the peptide ion population. D) Histogram of *m/z* of all peptides covered by the acquisition method in (C), and peptides not covered by the method but identified in a separately recorded spectral library. E) Number of peptide ions per isolation window. F) Histogram of ion mobilities of all peptides covered by the acquisition method, and peptides not covered by the method but identified in a separately recorded spectral library. The subfigures C-F are based on a reference proteome library (see Experimental Procedures).

Due to software constraints, the original dia-PASEF methods (6) comprise a repeating pattern of the top ion mobility windows per dia-PASEF scan. This leads to a configuration with equidistant quadrupole isolation widths (Fig. 1B). As a result, covering a wide *m/z* range comes at the cost of a high cycle time and reduced quantitative accuracy due to lower elution peak coverage. Alternatively, many peptide ions outside the *m/z* range would not be included in the acquisition scheme (Fig. 1C, D).

Moreover, when using equidistant isolation windows, the distribution of peptide ions per window is imbalanced, resulting in a high spectral complexity in highly dense regions (Fig. 1E). Lastly, this scheme for acquisition window setting is also suboptimal in the ion mobility dimension (Fig. 1F).

### Establishing an optimal dia-PASEF window design

We first investigated the optimum balance between the number of dia-PASEF scans and ion mobility windows per dia-PASEF scan to obtain a deep proteome coverage and quantitative accuracy. As described above, the original dia-PASEF method included three ion mobility windows per dia-PASEF scan. Having more ion mobility windows per dia-PASEF scan reduces cycle time but also diminishes precursor coverage due to smaller isolation windows in the ion mobility dimension (supplemental Fig. S5A, B). For instance, splitting the isolation width into two parts halved the complexity per spectrum and thereby increased identifications. However, doubling the number of dia-PASEF scans increases cycle time, which worsens the quantitative accuracy since only half as many data points are collected over one elution peak (supplemental Fig. S5A). We tested the impact of increasing the number of ion mobility windows per dia-PASEF scan and found that two ion mobility windows per dia-PASEF scan are optimal (supplemental Fig. S5C). Six points per elution peak are thought to be necessary for accurate quantitation (43). In the case of 21 minute gradients (60 SPD), we empirically found a median peak width of 8.27 s with our set-up (supplemental Fig. S5D) only considering precursors with a CV value below 20%. Each individual dia-PASEF scan takes around 100 ms plus one 100 ms MS1 scan per cycle and overhead time. Hence, 12 dia-PASEF scans amount to a cycle time of 1.38 s, ensuring adequate quantitative representation of the LC elution peak (see Experimental Procedures, supplemental Fig. S5E).

Given the limitations of our previous two-dimensional acquisition scheme, we needed to place and adjust *m/z* and ion mobility isolation windows flexibly. Existing tools such as “Define dia-PASEF Region” in Compass DataAnalysis (Bruker) or the “dia-PASEF window Editor” in TimsControl (Bruker) require the manual fitting of the scan area onto the peptide ion population and only generate isolation windows with equidistant widths. Therefore, we developed a Python package for Data Independent Acquisition with an Automated Isolation Design (py_diAID). It places two-dimensional dia-PASEF acquisition schemes in the *m/z*-ion mobility plane based on desired parameters (number of dia-PASEF scans, covered *m/z* and ion mobility range, and cycle time) and the empirical acquired reference data, which can be a proteomics library containing precursor ion information. The algorithms in py_diAID optimally adjust the variable quadrupole isolation widths according to the precursor density, aiming for an equal number of precursors fragmented per isolation window. Our simulations show that variable isolation widths enable short acquisition cycles covering essentially the entire *m/z*-ion mobility range (Fig. 2A, right panel).

**Figure 2:**
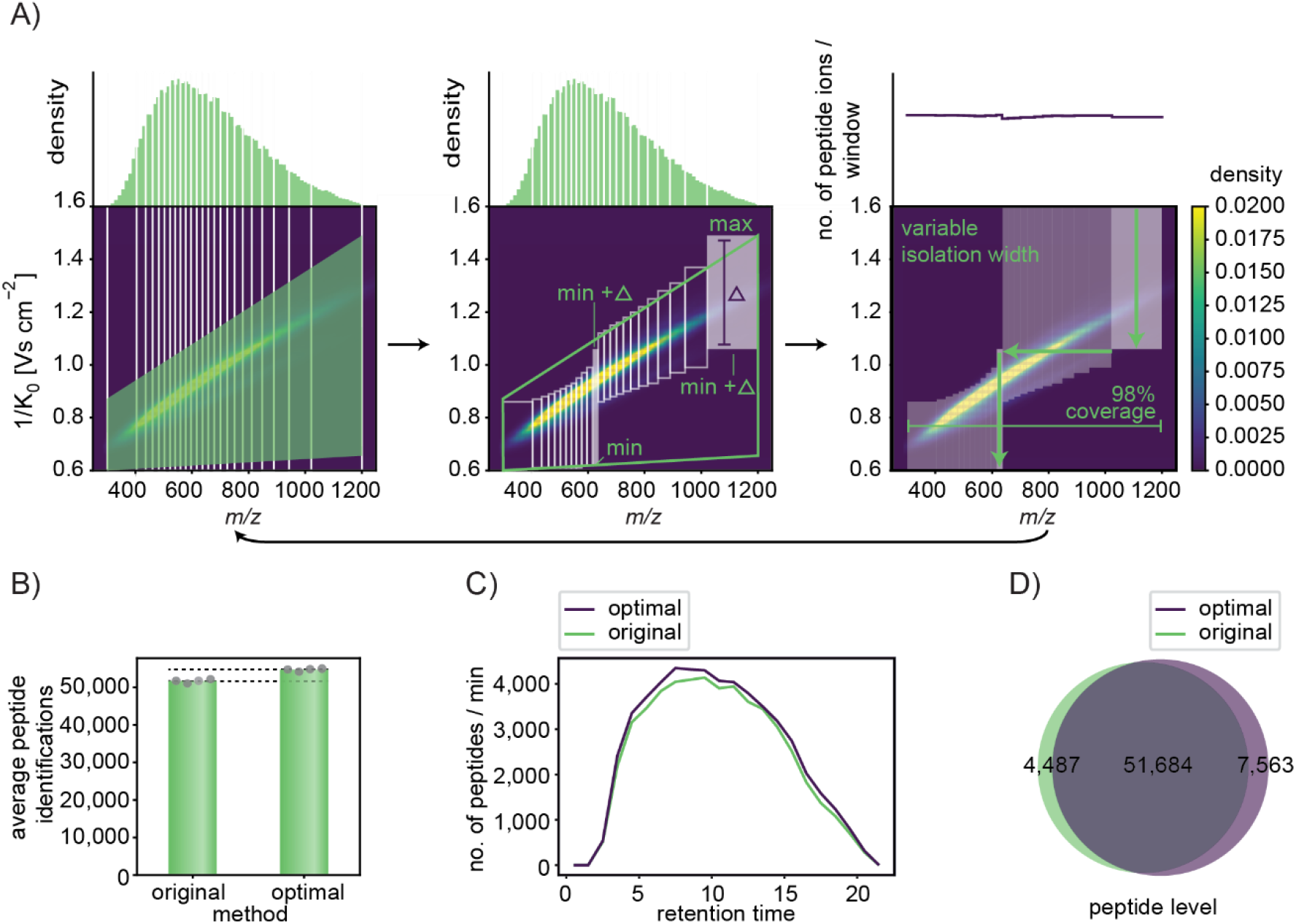
py_diAID algorithm and evaluation. A) py_diAID design of the optimal acquisition scheme and window placement for a 21 min gradient (60 SPD, Evosep) with variable widths to balance the distribution of peptide ions, providing nearly complete peptide ion coverage. The left panel illustrates the first steps of the py_diAID algorithm: defining the *m/z*-range of interest, binning the peptide ions in the *m/z*-dimension and definition of the scan area in the IM dimension. middle panel: Calculation of the isolation window dimensions and coordinates based on the scan area. right panel: Extension of the isolation windows to the limits of the ion mobility ranges. The arrow at the bottom indicates that the py_diAID algorithm evaluates the new acquisition scheme, defines the following test set of scan area parameters by Bayesian optimization, and resumes with the steps in the left panel. This is repeated for a user-defined number of iterations (more details in supplemental Fig. S6). A is plotted on top of a kernel density distribution based on the reference proteome library. B) Average peptide identifications by the original and optimal dia-PASEF methods. C) Number of peptides identified per minute over the entire retention time. D) Venn diagram showing the shared and unique peptides identified by both methods. Data in B to D are from quadruplicate injections of 200 ng tryptic HeLa digest with a 21 min gradient and analyzed with the reference proteome library.

Our algorithm first bins the precursor ion populations equally along the *m/z*-dimension. A trapezoid defines the extent of scan area and the position of the acquisition scheme in the *m/z*-ion mobility plane (Fig. 2A, left panel). Based on this, py_diAID calculates the optimal dimensions of each isolation window (Fig. 2A, middle panel) and extends the top and bottom ion mobility windows to the limits of the measured ion mobility range to maximize the covered peptide ion population (Fig. 2A, right panel and supplemental Fig. S6). The selected mass window of the quadrupole jumps at the determined transition point of each ion mobility window within each dia-PASEF scan. In each subsequent dia-PASEF scan, the starting *m/z* window is offset to lower values based on the individual width of the previous window (Fig. 2A). Next, py_diAID evaluates the generated acquisition scheme based on the covered precursor ions of an experimentally acquired library or subset thereof, for example one filtered by a charge state or by a population of modified peptides. This is a multivariant non-linear optimization problem, and we used the gp_minimize module provided by the Scikit-Optimize (skopt) library in Python to perform this task that is highly used in machine and deep learning for the hyperparameter optimization (see Experimental Procedures). Its inputs are the trapezoid corners and it iteratively decides which parameters should be tested next based on the above evaluation. This process is repeated for many iterations (about 200 in practice, supplemental Fig. S7) until it converges to the best window placement. py_diAID is available as a Python module, a command-line interface, and a graphical user interface on all major operating systems under an Apache 2.0 license (supplemental Fig. S8). The source code is freely available on GitHub (https://github.com/MannLabs/pydiAID).

We first benchmarked the optimal dia-PASEF methods designed with py_diAID against the original dia-PASEF method. The optimal dia-PASEF method calculated by py_diAID covered 99% of all doubly and 94% of all triply charged precursors in the ‘reference library’, that was generated with FragPipe. This compares to 88% and 71% coverage with the original dia-PASEF. The original dia-PASEF method had already been extensively and manually optimized for the short gradient lengths and the tryptic HeLa digest employed here. This explains why the number of experimentally identified proteins is very similar between both methods (supplemental Table 1). However, even in this case, py_diAID’s optimal acquisition scheme increased the number of identified peptides by 6% in single-run injections (Fig. 2B) and across the entire retention time (Fig. 2C), while the number of peptide identifications in replicate injections deviates only by 1%. Inspection of the data shows that the additional peptides originate both from the previously not covered regions and from the most dense elution times. More than 80% of all identified peptides were commonly identified by both methods (Fig. 2D). In other applications, such as phosphoproteomics, the gains by py_diAID were much larger (see below Fig. 5).

### Deep proteome coverage in short LC-gradients

We next investigated if coupling our optimized dia-PASEF methods with project-specific, in-depth libraries yields higher peptide identification and improves quantification accuracy. To generate such an in-depth library, we separated 15 μg of the HeLa sample that we also used for single dia-PASEF acquisitions into 48 concatenated fractions using high pH reverse phase chromatography of the Spider fractionator (see Experimental Procedures) (31). These fractions were measured in dda-PASEF mode and again analyzed with FragPipe and its SpecLib workflow. We compared our ‘reference library’ generated with limited sample amount (2.5 μg proteolytic digest) and 24 fractions to the new one with ample sample amount (15 μg) and twice as many fractions. As expected, the latter was substantially larger, containing 45% more peptides (counting all modifications) and 13% more proteins. Altogether, this deep library constructed from 21 min runs comprised 124,155 peptides and 8,439 different protein groups (Fig. 3A, B).

**Figure 3:**
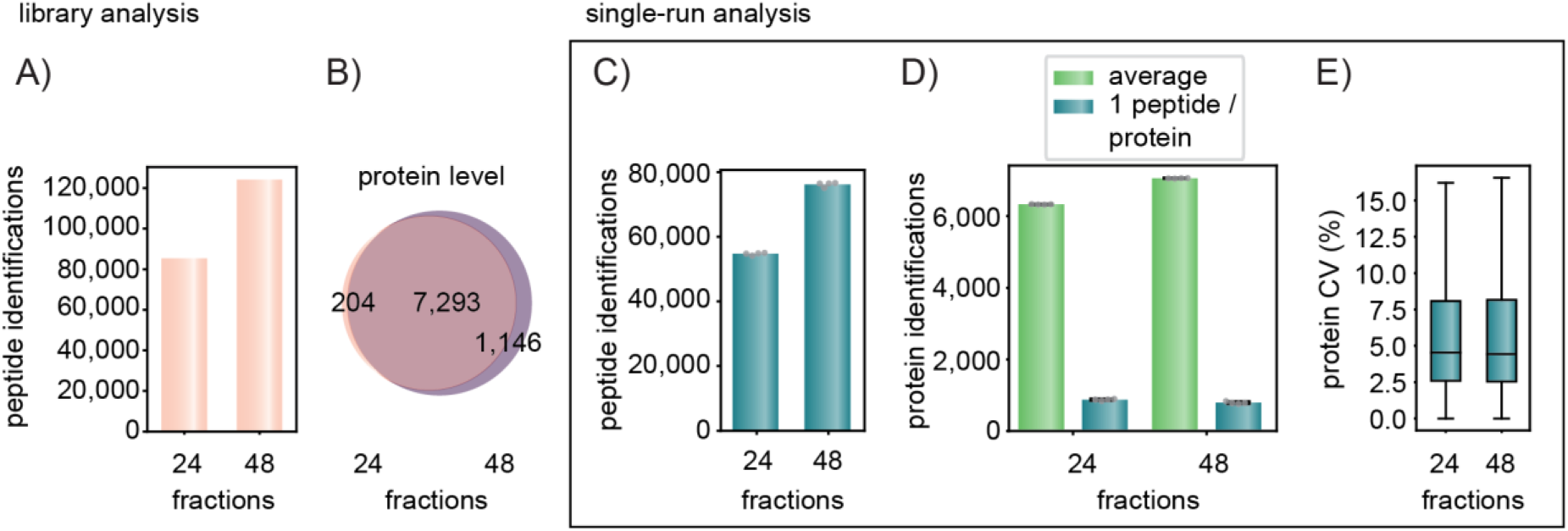
Workflow optimization for the 21 min gradient with project-specific deep libraries. A) Peptides identified of the reference vs. the project-specific deep library for 21 min runs. B) Shared proteins and depth on the protein level in the two libraries. C) Average peptide identification of four single-run injections. This data and the one in (D) and (E) were generated from quadruplicate injections of 200 ng tryptic HeLa digest acquired with a 21 min gradient and searched with the reference (24 fractions) or project-specific library (48 fractions). D) Average protein identifications and identifications with only one peptide in the single runs. E) Coefficients of variation at the protein level based on the MaxLFQ algorithm of DIA-NN. Boxplots show the median (center line), 25^th^, and 75th percentiles (lower and upper box limits, respectively), and the 1.5× interquartile range (whiskers). n = 6,384 (24 fractions) and 7,121 (48 fractions) shown in panel C.

Next, we compared single dia-PASEF runs with reference vs deep library using DIA-NN and found a corresponding increase in the proteome depth (39% more peptides and 12% more proteins) (Fig. 3C, D). Using the deep library identified 76,214 ± 1,021 peptides and the reference library 51,711 ± 641 peptides (Fig. 3C). With the deep library, an astounding 7,056 ± 8 proteins were identified with our optimized acquisition scheme in each of four replicate runs on average. Specifically, with the reference library, DIA-NN reported 14% significant protein identifications on the basis of one peptide and this percentage decreased slightly to 11% with the deeper library (Fig. 3D).

Quantitative reproducibility between the quadruplicates was virtually identical when using the reference or deep library (4.5% vs 4.4% on protein level and 12.1% vs 13.45% on peptide level) (Fig. 3E). Taken together, we found that single run identification benefited from a project-specific, in-depth library while maintaining the accuracy of quantification. We therefore used the 48 fractions library for all 21 min runs to generate equivalent libraries for evaluating a range of gradient lengths as described next (referred to as ‘project-specific deep libraries’).

We next investigated the effect of even shorter gradients as well as somewhat longer gradients on proteome depths and quantitative accuracy. As before, each library was acquired with dda-PASEF and 15 μg HeLa lysate separated into 48 fractions. Extending the gradient to 44 min (30 SPD method on the Evosep One system) identified an average of 7,756 ± 6 proteins based on 100,900 ± 634 peptides (including all modifications). This represents an identification increase of 10% on protein level in comparison to the 21 minutes gradient. The median CV between the quadruplicates was 4% at the protein level for these technical replicates, and 7,393 protein groups had CVs below 20% (Fig. 4A, C).

**Figure 4:**
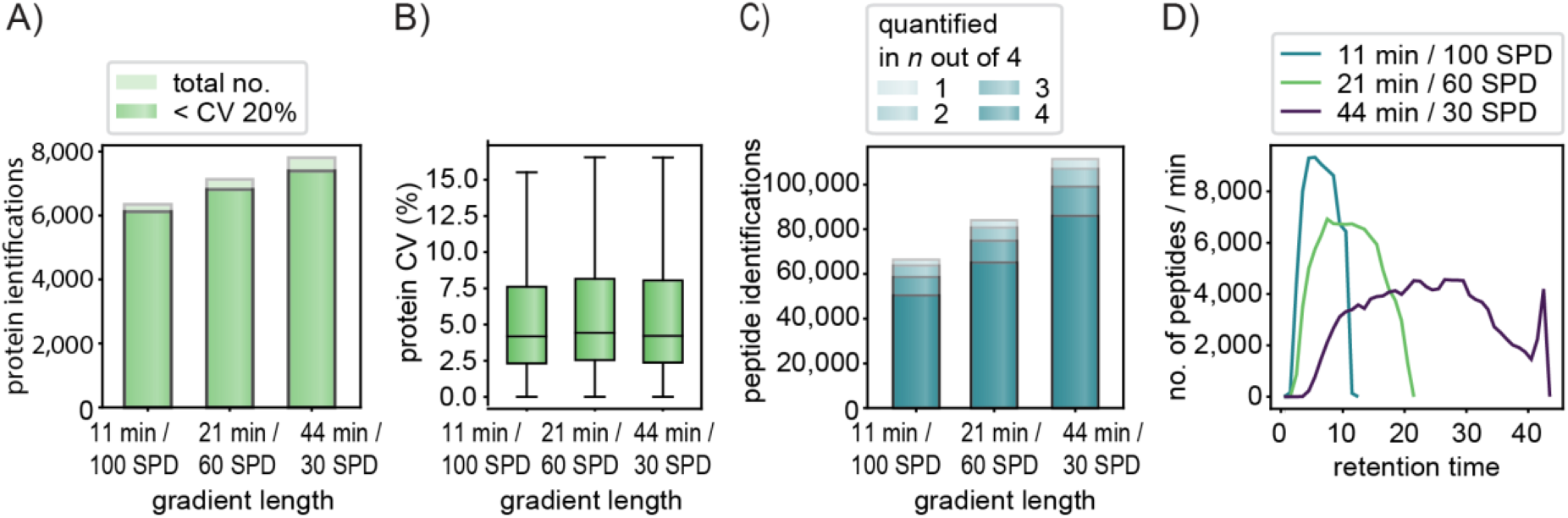
Comparison of different gradient lengths/ throughput based on single-run analysis. A) All single-run identifications and those with a CV < 20% for the 11 min, 21 min and 44 min gradients. B) Coefficients of variation at the protein level based on the MaxLFQ algorithm of DIA-NN. Boxplots show the median (center line), 25^th^, and 75th percentiles (lower and upper box limits, respectively), and the 1.5× interquartile range (whiskers). n = 6,341 (11min / 100 SPD) and 7,121 (21min / 60 SPD), and 7,802 (44min / 30 SPD) shown in panel A. C) Analysis of peptide quantification in n out of four technical replicates shows that the large majority is quantified consistently. D) The number of peptides per second over the retention time for the three gradient lengths. The data was acquired in quadruplicate injections of 200 ng HeLa digest and analyzed with 48 fraction, dda-PASEF libraries each recorded with the corresponding gradient length. 11-min library: 8,553 proteins and 122,105 peptides, 21-min library: 8,439 proteins and 124,155 peptides, 44-min library: 9,461 proteins, 175,839 peptides.

**Figure 5:**
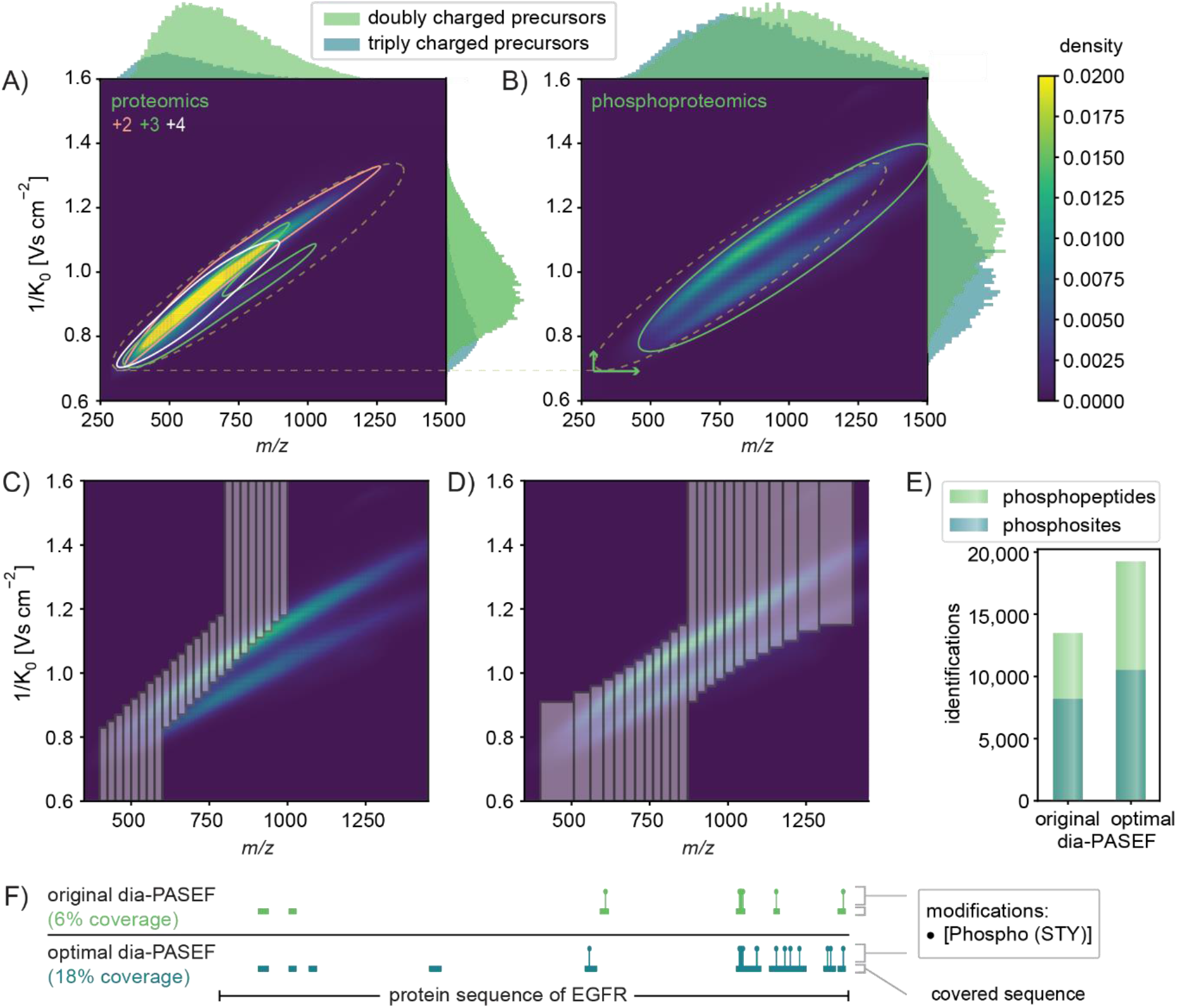
Method optimization specifically for phosphoproteomics. A) Peptide distribution of a proteomics digest displayed as kernel density estimation dependent on the charge and histograms of the abundance of differently charged precursors based on our deep proteomics library. B) Peptide distribution of a phosphoproteomics digest displayed as kernel density estimation and histograms of the abundance of differently charged precursors based on our phosphopeptide library. C) Original dia-PASEF method plotted on top of the phosphopeptide library. D) Optimal dia-PASEF method tailored to the phospho-library. E) Identified phosphosites and phosphopeptides based on quadruplicates of 100 μg EGF-stimulated and enriched HeLa digest, separated within 21 min and searched with DIA-NN against the phospho-library. F) AlphaMap visualization (48): Protein sequence coverage of the epidermal growth factor receptor (EGFR) depending on the acquisition method.

We expected that the fast scan rate of the timsTOF, together with our optimized method might still accurately measure a large part of the proteome even in very short gradients (6, 32). Indeed, the 100 SPD method (11 min gradient) still identified 6,285 ± 18 proteins (59,811 ± 368 peptides). Quantitative accuracy reported by DIA-NN did not suffer and remained at a median CV of 4%. Taking only the proteins with CVs equal or below 20%, the 100 SPD method still resulted in 6,121 proteins, covering 83% of proteins that could be accurately quantified with the 44 min gradient while substantially reducing the analysis time (Fig. 4A). Rank order reproducibility was also high for these technical replicates for all gradient lengths (supplemental Fig. S9-10, r=0.999 for proteins and r=0.992 for peptides). As expected, the number of peptides identified per minute decreased when increasing the gradient length while the 11-min gradient reached the highest numbers (9,330 peptides per minute translating to 155 peptide identifications per second at the apex, Fig. 4D).

In conclusion, our data show that our improved workflow constitutes a powerful technological platform capable of accurately quantifying a large part of the proteome at high throughput.

### Comparison of proteome results between DIA-NN and Spectronaut

The above analyses were all performed with the DIA-NN package. To determine if our results depend on the software used, we employed Spectronaut (Biognosys) (3), another widely used software package (11, 44). This revealed that both packages identified comparable numbers of proteins. For instance, in the 60 samples per day method, Spectronaut reported 7,285 significant protein groups, whereas DIA-NN reported 7,056 (supplemental Fig. S11 A). In the version tested (Spectronaut 16), this also held for even shorter gradients (6,250 vs. 6,285). Having established that the overall protein numbers are similar, we next investigated the overlap between the found proteins. As DIA-NN has a different protein grouping algorithm from Spectronaut, we performed this analysis on the level of genes and peptide precursors. Employing similar grouping schemes at the gene level showed a high level of concordance, with 548 genes unique to Spectronaut and 208 unique to DIA-NN out of a total of 7,668 identified genes for both (supplemental Fig. S11 B). For the total of 128,002 identified peptide precursors, the discrepancy was somewhat larger, with 28% unique identifications for Spectronaut and 5% for DIA-NN (supplemental Fig. S11 C). Overall, based on these proteome results, we conclude that the gains achieved by py_diAID are independent of the DIA analysis software used.

### Rapid phosphoproteomics with optimal isolation window design

Phosphorylation, one of the most prevalent and most studied post-translational modification, refers to the addition of a phosphoryl group – usually on serine, threonine or tyrosine amino acid residues. This introduces a mass and ion mobility shift on the modified peptides, indicating that analysis of phosphopeptides can benefit from the additional ion mobility dimension in PASEF (45, 46). To date, dia-PASEF has not been explored in a large-scale study of the phosphoproteome or any other post-translationally modified sub-proteome.

It is well known that the ion mobility dimension separates peptides in clouds primarily reflecting their charge status. In the timsTOF case, Figure 5A depicts dense clouds containing doubly, triply, and quadruply charged peptide ions (47). In the case of phospho-enriched samples, projecting the distribution of phosphorylated peptides into the *m/z* and IM space revealed a substantial shift of ion cloud to higher *m/z* values and higher IM values, due to the 80 Da increase in their mass, higher charge states and conformational changes upon phosphorylation (Fig. 5B). These observations suggest that dia-PASEF methods need to be tailored for phosphoproteomics. To this end, we first generated an in-depth phospho-library from EGF stimulated HeLa cells that were separated into 76 fractions and then enriched for phosphorylated peptides. These enriched fractions were measured with the 60 SPD method, dda-PASEF in little more than one day. We analyzed the results both by FragPipe combined with DIA-NN and by Spectronaut 16 (see Experimental Procedures). This generated an in-depth library of 187,730 modified or unmodified peptides, 123,133 phosphopeptides and 107,154 phosphosites for DIA-NN. Spectronaut 16 obtained very similar results (194,309 modified or unmodified peptides, 132,270 phosphopeptides and 114,158 phosphosites). The overlap between phosphopeptides was 50% (supplemental Fig. S13A).

When we simulated the coverage of the original dia-PASEF method for the 21 min gradient (6), we found that it only reached a coverage of 34% of phosphopeptide ions in our deep phospho-library, in contrast to the 81% achieved for unmodified peptides (Fig. 5C). Therefore, we used our phospho-library as input for py_diAID to obtain a dia-PASEF method tailored for phosphoproteomics. This resulted in a theoretical coverage of 93% of all doubly charged and 92% of all triply charged phosphopeptide ions (Fig. 5D).

We next utilized this optimal dia-PASEF phospho-method to measure the samples containing phosphorylated peptides enriched from 100 μg digest of EGF stimulated HeLa cells. We first analyzed the resulting files with DIA-NN against our deep phospho-library. In agreement with our simulations, the original dia-PASEF method identified 8,199 phosphosites and 13,485 phosphopeptides whereas the optimal method detected 28% more phosphosites (10,510) and 43% more phosphorylated peptides (19,258) (Fig. 5E). To illustrate this further, we mapped the experimentally acquired phosphopeptides to the EGF receptor (EGFR) sequence essential for transmitting the EGF signal using AlphaMap (48). This revealed that the optimal dia-PASEF phospho-method doubled the number of detected phosphosites to a total of 14 (Fig. 5F).

The intensities of the phosphopeptides detected in our deep FragPipe phospho-library in dda-PASEF mode and 76 fractions span almost seven orders of magnitude (supplemental Fig. S12A). When searching single dia-PASEF phospho-runs against our phospho-library using DIA-NN, we found that single short gradients covered 21% of the phosphopeptide sequences, ranging from 12% in the most abundant quintile to 0.3% in the least abundant one (Suppl. Fig. S12A). Apart from the statistical analysis, the AlphaViz package (49), based on AlphaTims (50), allows visualization of any phosphopeptides of interest. This is shown for the phosphopeptide ELVEPLT[Phospho (STY)]PSGEAPNQALLR on EGFR, where the distinct precursor and fragment peaks are clearly visible in the retention time dimension and even more important in the retention time – ion mobility plane, supporting the DIA-NN assignment (supplemental Fig. S12B, C).

Next, we analyzed the same single-run phospho dataset with Spectronaut. To our surprise – especially given the comparable results at the proteome level – Spectronaut drastically increased the number of identified phosphosites to 28,980 (supplemental Fig. S13B). This was even more pronounced for identified phospho-peptides (72,216, supplemental Fig. S13C). Accordingly, the common overlap of phosphosites was only 26% (supplemental Fig. S13B). We do not know the origin of this large discrepancy, but we encourage the providers of these software packages to resolve this, especially as the code is not available for inspection. In the context of our study, we decided to continue with the more extensive Spectronaut results, as they appeared to still correctly represent the regulation in the EGFR signaling experiment described below.

### In-depth phosphoproteomics analysis of the EGF-signaling pathway

To benchmark our optimal dia-PASEF workflow, we chose the well-studied epidermal growth factor (EGF) signaling pathway in HeLa cells. The binding of EGF to the EGF receptor (EGFR) results in the activation of downstream kinases, which phosphorylate a repertoire of numerous substrates, regulating diverse cellular processes (51). We aimed to quantitatively and accurately measure the differential phosphorylation of proteins involved in this signaling pathway using our rapid and sensitive method. To this end, EGF-treated and control samples were collected in three biological replicates, digested into peptides and enriched for phosphorylated peptides (see Experimental Procedures). Subsequently, we measured the enriched phosphopeptides with dia-PASEF in 21 minutes and searched the deep phosphopeptide library that we already employed for the method optimization described above with Spectronaut 16.

With our workflow, we quantified 46,136 phosphorylation sites on 4,300 proteins. Of these, 35,537 sites were identified with a high site localization probability (75%, Class I sites (52)) and 20,001 were quantified in all replicates of at least one experimental condition (Fig. 6A). The dia-PASEF workflow allowed high reproducible quantification demonstrated by a median Pearson coefficient above 0.92 for replicates within conditions (Fig. 6B). Remarkably, a full 26% (5,200, 5% FDR) and 10.5% (2,117, 1% FDR) of phosphorylation sites were significantly modulated upon EGF treatment (Fig. 6C).

**Figure 6:**
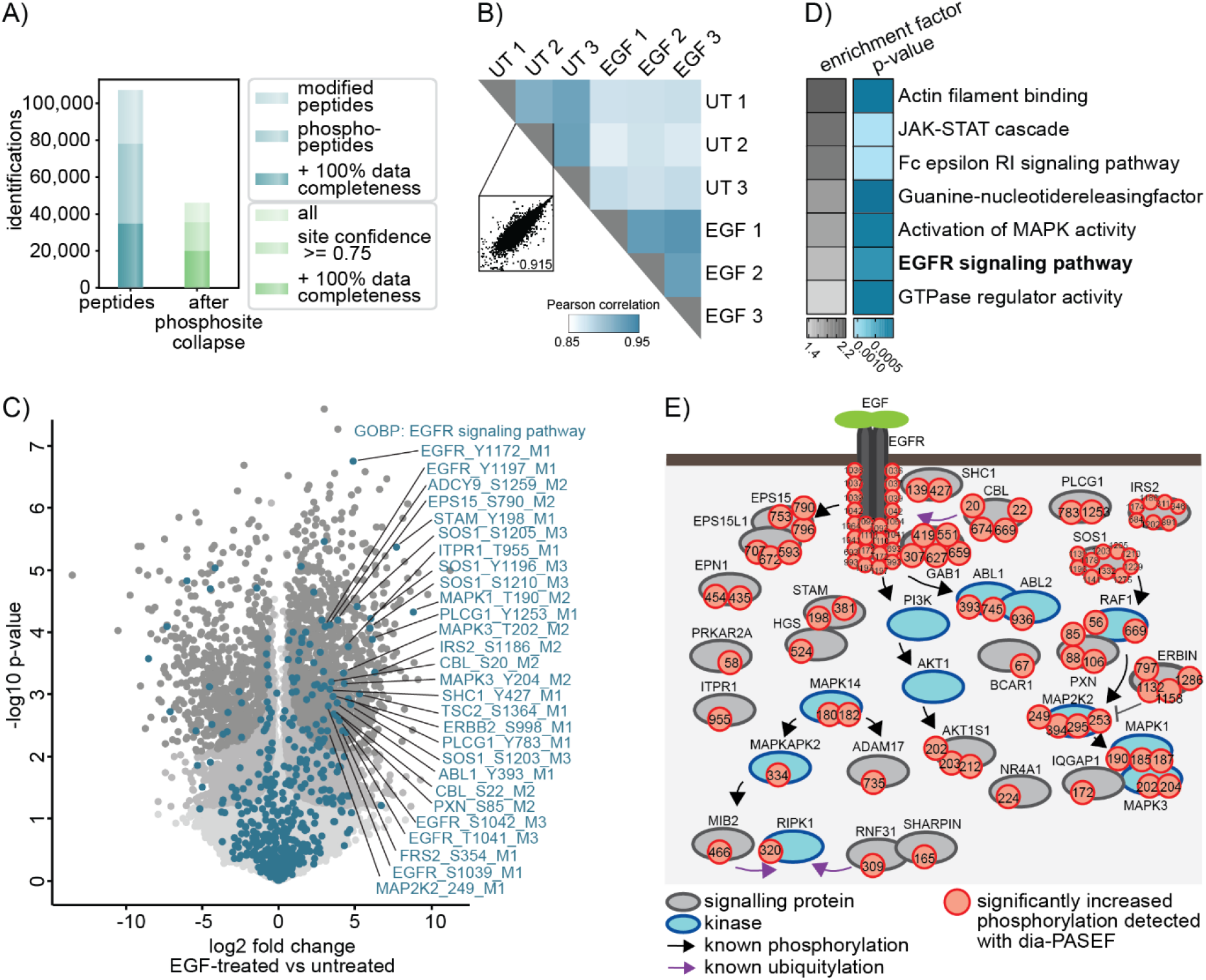
The dia-PASEF workflow allows the robust detection of characteristic EGF signaling events. A) Numbers of all identified phosphopeptides and phosphosites before and after filtering for localization probability and data completeness. B) Phosphoproteome Pearson correlation matrix. Scatter plot shows the correlation of replicates within a condition. C) Volcano plot of phosphosites regulated upon 15 min of EGF-treatment in Hela cells vs untreated cells. (two-sided Student’s t-test, FDR<0.1 = grey, FDR<0.5 = dark grey). Protein’s part of the GOBP term ‘EGFR signaling pathway’ are highlighted in turquoise. D) Fisher’s exact test of proteins with significantly increased phosphosites upon EGF treatment (p-value<0.002). Enrichment annotations are GOBP, GOMF and KEGG. E) Scheme of significantly upregulated phosphosites that were detected in this study and are part of the GOBP term ‘EGFR signaling pathway’ and/or changed significantly upon EGF stimulation (FDR<0.05).

As expected, Gene Ontology (GO) enrichment analysis revealed strong overrepresentation of proteins involved in the EGFR signaling pathway (GOBP) and related pathways among the significantly EGF-upregulated phosphoproteins (Fig. 6D). Most are known to be critical for intact EGF signaling. For example, we detected phosphorylation of T693, Y1110, Y1172, Y1197 on the receptor EGFR itself, Y427 on the adaptor protein Src Homology 2 Domain-Containing-Transforming Protein C1 (SHC1), Y659 on Growth Factor Receptor Bound Protein 2-Associated Protein 1 (GAB1) and on the downstream kinases Mitogen-Activated Protein Kinase 2 (MAP2K2) (T394) and Mitogen-Activated Protein Kinase 1 and 3 (MAPK1/3) (T185/Y187, T202/Y204) (53) (Fig. 6C, E). These phosphosites are typically used to examine EGF signaling with classical methods such as immunoblotting or with targeted mass spectrometry (9, 10, 54). These approaches, however, only allow relatively low throughput analyses, that require dedicated assay development procedures or the generation of phosphospecific antibodies. In contrast, by combining the automated phosphoenrichment on the BRAVO platform with the robust Evosep and timsTOF setup, our approach achieves 60 SPD. This allows us to track and accurately quantify the induction of more than 60 phosphorylation events on proteins critical for EGF signaling (part of GOBP: EGFR signaling pathway) within a single 21-min run (supplemental Fig. S14). Importantly, besides the phosphorylations of the classical EGF signaling members, many other signaling events that, for example, result from signaling crosstalk downstream of the EGF receptor can also be detected, including S897 of the Ephrin Type-A Receptor 2 (EPHA2), S339 of the C-X-C Motif Chemokine Receptor 4 (CXCR4) and T701 of Erb-B2 Receptor Tyrosine Kinase 2 (ERBB2) (supplemental Fig. S14).

To identify functionally important phosphorylation events not directly linked to EGF signaling, we matched the functionality prediction score developed by Beltrao and co-workers to the upregulated phosphorylation events (55). We identified 659 phosphosites with a high functional score of >0.5 to be significantly upregulated, which are not part of the GOBP term ‘EGFR signaling pathway’ (FDR<0.05) (supplemental data 1). These include EGF-induced phosphorylation of E3 ligases like Mindbomb Homolog 2 (MIB2) (S309) and members of the linear ubiquitin chain assembly complex Ring Finger Protein 31 (RNF31) (S466) and Sharpin (S165), which are most frequently studied in the context of TNF signaling (supplemental Fig. S14) (56–59). Similarly, phosphorylation of Receptor Interacting Serine/Threonine Protein Kinase 1 (RIPK1) on S320, which prevents TNF-induced cell death, was also increased upon EGF signaling (supplemental Fig. S14) (60, 61). This phosphorylation is mediated by MAP kinase-Activated Protein Kinase 2 (MAPKAPK2), which is activated upon EGF stimulation demonstrated by its increased phosphorylation at T334. These are just some examples of functional candidates whose role in EGF signaling has still to be determined.

Together, this EGF study demonstrates the quantitative capabilities of the dia-PASEF-based phosphoproteomics workflow. We conclude that efficient analysis of ions separated in the IM and *m/z* space enables the investigation of signaling pathways with high sensitivity in a high-throughput manner.

## DISCUSSION

The optimal placement of dia-PASEF windows in the two-dimensional *m/z* and ion mobility space is not trivial. We here developed py_diAID which is available on GitHub at MannLabs and is installable as a Python module with a command line interface or as a GUI on Windows, Mac and Linux. It adjusts the isolation window width to the precursor density, and optimally positions the isolation design in the *m/z*-IM space. This leads to near-complete theoretical precursor coverage for proteomics. Compared to the original dia-PASEF method (6), the gains for phosphorylated precursors are especially striking (34% vs. 93%).

MS-based proteomics is a rapidly developing technology. For perspective, to cover ten thousand proteins we had to measure the samples for twelve days with four-hour gradients ten years ago (62). Here, we coupled a robust, high throughput LC system to the TIMS-qTOF instrument employing the rapid sampling speed of a TOF analyzer. It offers short gradients and also low overhead time, enhancing the overall throughput capabilities (27). With this, we generated in-depth project-specific libraries of 9,461 proteins in only 13% of the previous measurement time. Furthermore, once the libraries are ready, subsequent proteome characterization using py_diAID generated methods happens in only 44 minutes to a depth of 7,700 proteins (less than 1% of the measurement time necessary ten years ago). Our workflow is also twice as fast as currently employed high throughput screening strategies for cancer proteomics, while achieving greater proteome depth on cell lysate (63–65).

So far, there have been only a few reports of the timsTOF principle on phosphoproteomics (28). Here, we show that this instrument is capable of in-depth phosphoproteomics with very high sensitivity. Specifically, we identified thirty-five thousand phosphosites in only 21 minutes in triplicates from 100 μg EGF-stimulated HeLa cell digests. Our workflow opens up the possibility to measure multiple pathways in a short time. We demonstrated that quantitatively analyzing the regulated phosphoproteome covers the well-studied EGF signaling pathway together with auxiliary pathways. Interestingly, our workflow is even faster than selected reaction monitoring employed as a targeted screening method for assessing the activation of signaling pathways (9). However, our method is generic to any pathway and applicable in principle to the entire phosphoproteome.

In the current implementation, the dia-PASEF windows are adjusted based on empirical data before the acquisition. These adjustments could also be implemented in real-time based on the precursor density achieving an acquisition design optimized to the individual time points of an entire gradient. Furthermore, we employed in-depth libraries. While they can be generated quickly, current developments of *in silico* generated DIA libraries or direct DIA methods may soon obviate the need for this step. Likewise, we expect that py_diAID will perform similarly for other PTMs.

## ABBREVIATIONS

ABC: ammonium bicarbonate
CAN: acetonitrile
CAA: 2-chloroacetamide
Dda: data-dependent acquisition
Dia: data-independent acquisition
EGF: epidermal growth factor
FA: formic acid
GO: Gene Ontology
IM: ion mobility
IPA: isopropyl alcohol
KEGG: Kyoto Encyclopedia of Genes and Genomes
MeOH: methanol
PASEF: parallel accumulation – serial fragmentation
PBS: phosphate-buffered saline
PTM: post-translational modification
py_diAID: Python package for Data-Independent Acquisition with an Automated Isolation Design
SDC: sodium deoxycholate
SPD: samples per day
TBS: tris-buffered saline
TCEP: tris(2-carboxy(ethyl)phosphine)
TFA: trifluoroacetic acid
TIMS: trapped ion mobility spectrometry

## ACKNOWLEDGEMENT

We thank Nagarjuna Nagaraj for introducing us to the acquisition software timsControl, which is a basis for py_diAID, and our colleagues in the Department of Proteomics and Signal Transduction at the Max Planck Institute of Biochemistry. We are particularly grateful for the help from Ankit Sinha, Igor Paron, Maria Wahle, Corazon Ericka Mae Itang, Isabell Bludau, Constantin Ammar and Medini Steger.

## FUNDING AND ADDITIONAL INFORMATION

This study was supported by the Max-Planck Society for Advancement of Science, the Deutsche Forschungsgemeinschaft (DFG) project ‘Chemical proteomics inside us’ (grant 412136960) and by the Bavarian State Ministry of Health and Care through the research project DigiMed Bayern (www.digimed-bayern.de).

## DATA AVAILABILITY

All dia-PASEF parameter files required for the acquisition, mass spectrometry raw files corresponding to the spectral libraries and single-run experiments, and output information from DIA-NN and MS-Fragger have been deposited with the ProteomeXchange Consortium *via* the PRIDE partner (66) repository with the dataset identifier PXD034128. Supplemental data 2 is a roadmap linking the raw files. Homo sapiens (taxon identifier: 9606) proteome databases were downloaded from https://www.uniprot.org. py_diAID is a fully open-source package, and the code is freely available under the Apache 2.0 license at https://github.com/MannLabs/pydiAID.

## AUTHOR CONTRIBUTIONS

P.S., M.T., M.C.T., F.M.H., Ö.K., A.-D.B., F.M and M.M. conceptualized and designed the study; P.S. and F.M. conceived the tool py_diAID; P.S., M.T., M.C.T. and F.M.H. performed experiments; P.S., E.V. and S.W. developed the tool py_diAID; P.S., M.T., M.C.T., E.V., F.M.H., Ö.K., A.-D.B., F.M and M.M. analyzed the data; P.S., M.T., S.W., Ö.K. and M.M. wrote the manuscript with input from all authors.

## CONFLICT OF INTEREST

M. M. is an indirect investor in Evosep Biosystems. All other authors declare that they have no conflicts of interest with the contents of this article.

## SUPPLEMENTAL DATA

This article contains supplemental data.

## Notes

https://github.com/MannLabs/pydiaid

