## supplementary information for "Rapid and in-depth coverage of the (phospho-)proteome with deep libraries and optimal window design for dia-PASEF"

### SUPPLEMENT

**Supplementary Table 1: Pooled fractions for generating the phosphoproteomics library. All other fractions were acquired as individual runs.**

| Pooled fractions |
|---|
| 1, 90-95 |
| 2, 73 |
| 74, 75, 82, 83 |
| 76, 77, 84, 85 |
| 78, 79, 86, 87 |
| 80, 81, 88, 89 |

**Supplementary Table 2: Proteomics libraries.**

| Library specificities | No. of fractions / analysis software | No. of precursors | No. of modified peptides | No. of peptides | No. of proteins |
|---|---|---|---|---|---|
| 11 min / 100 SPD | 48 / FragPipe | 147,525 | 122,105 | 99,138 | 8,553 |
| 21 min / 60 SPD | 24 / FragPipe | 99,737 | 85,374 | 74,323 | 7,497 |
| 21min / 60 SPD | 48 / FragPipe | 148,244 | 124,155 | 98,061 | 8,439 |
| 44 min / 60 SPD | 48/ FragPipe | 219,736 | 175,839 | 137,582 | 9,461 |
| 11 min / 100 SPD | 48 / Spectronaut | 139,673 | 115,015 | 99,962 | 8,583 |
| 21min / 60 SPD | 48 / Spectronaut | 139,198 | 116,407 | 98,623 | 8,519 |

**Supplementary Table 3: Identifications on protein level.**

| | original dia-PASEF | optimal dia-PASEF |
|---|---|---|
| protein groups with CV <10% (MaxLFQ) | 5284 | 5271 |
| protein groups with CV <20% (MaxLFQ) | 6188 | 6155 |
| average identifications | 6337.25 | 6328.25 |
| total identifications | 6395 | 6386 |

**Supplementary Data 1: Phosphoproteome analysis of EGF-treated HeLa cells.**
This file reports the fold changes, the significance (q-value, two-sided Student's t-test), the uniport IDs, the PTM collapse key, and the functionality score of phosphosites that are significantly upregulated (FDR < 0.05) upon EGF stimulation and have a functionality score > 0.5 (55), but are not part of the GOBP 'EGFR signaling pathway'.

**Supplementary Data 2: Raw files used for analysis.**
This file links raw files to their dia-PASEF method, library, analysis output tables, post-analysis tables, and figures.

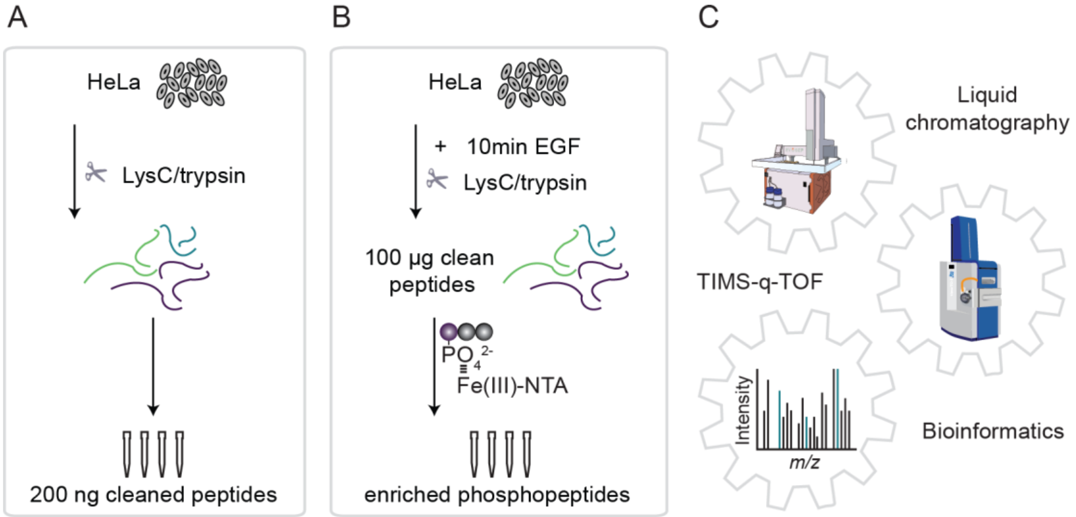


**Supplementary Figure S1:**
A) Experimental set-up for proteomics experiments in quadruplicates.
B) Experimental set-up for phosphoproteomics experiments in quadruplicates.
C) LC-MS set-up and data analysis with bioanalytic software (DIA-NN and, for comparison, Spectronaut).

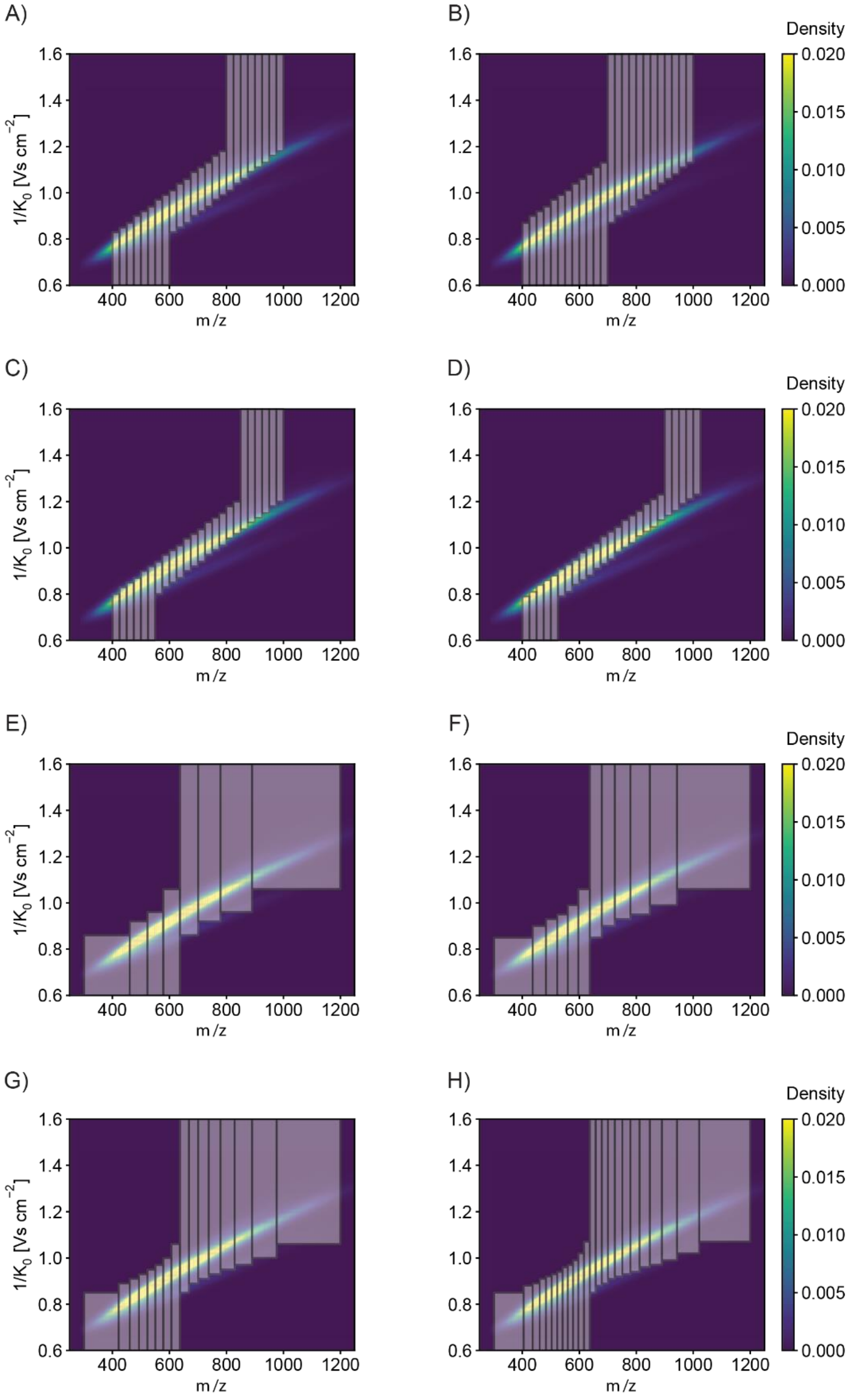


**Supplementary Figure S2: dia-PASEF method plotted on top of kernel density estimation of 'reference proteome library'. Duty cycle = 1 / No. of dia-PASEF scans. The duty cycle indicates the used proportion of the ion beam.**

A) Original dia-PASEF method (6), 12.5% duty cycle.
B) dia-PASEF method with 2 IM windows / dia-PASEF scan, 8.3% duty cycle.
C) dia-PASEF method with 4 IM windows / dia-PASEF scan, 16.7% duty cycle.
D) dia-PASEF method with 5 IM windows / dia-PASEF scan, 20% duty cycle.
E) dia-PASEF method with 4 dia-PASEF scans, 25% duty cycle.
F) dia-PASEF method with 6 dia-PASEF scans, 16.7% duty cycle.
G) dia-PASEF method with 8 dia-PASEF scans, 12.5% duty cycle.
H) dia-PASEF method with 12 dia-PASEF scans, 8.3% duty cycle.

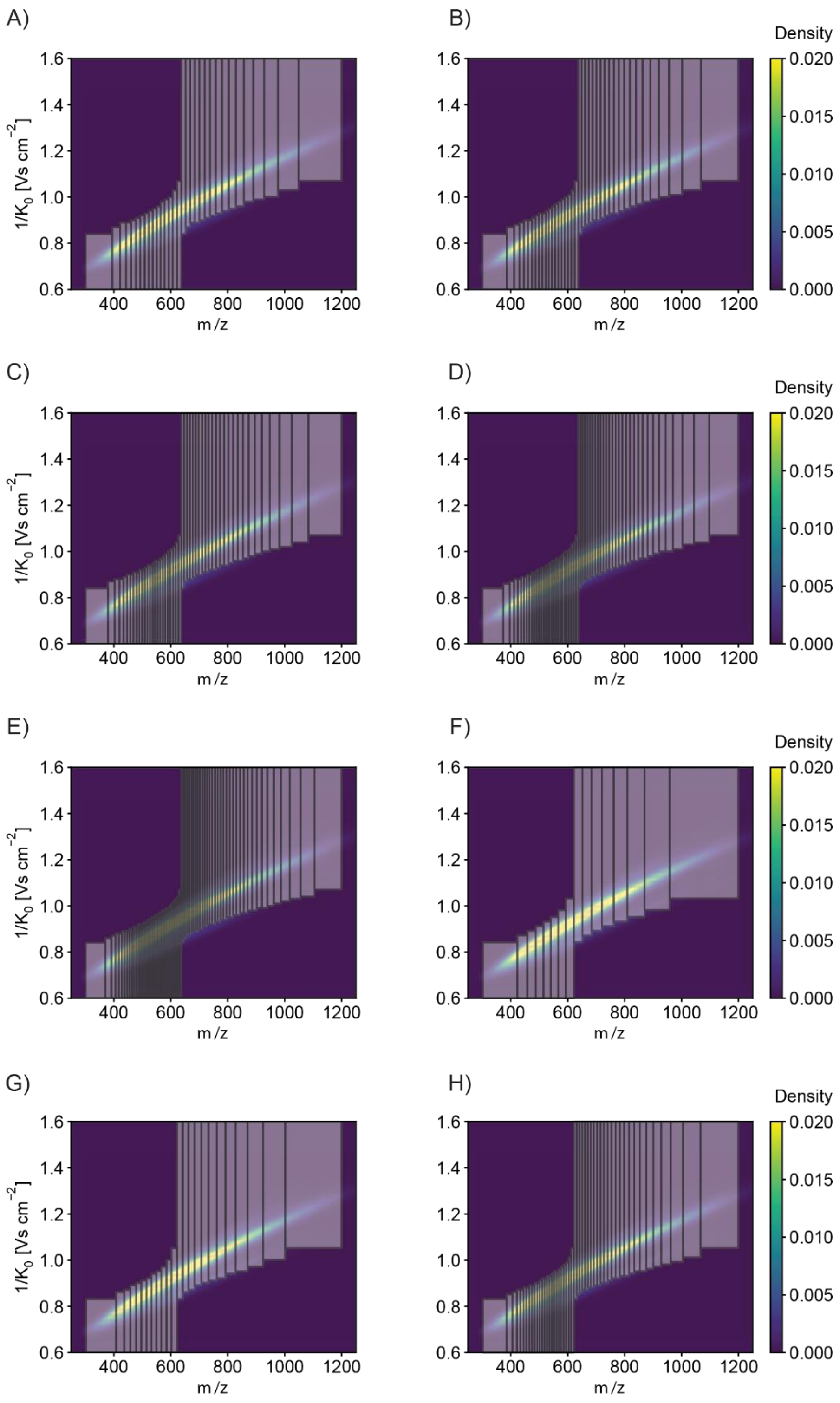


**Supplementary Figure S3: dia-PASEF method plotted on top of kernel density estimation of 'reference proteome library'.**

A) dia-PASEF method with 16 dia-PASEF scans, 6.3% duty cycle.
B) dia-PASEF method with 20 dia-PASEF scans, 5% duty cycle.
C) dia-PASEF method with 25 dia-PASEF scans, 4% duty cycle.
D) dia-PASEF method with 30 dia-PASEF scans, 3.3% duty cycle.
E) dia-PASEF method with 35 dia-PASEF scans, 2.9% duty cycle.
F) dia-PASEF method for 11 min gradients / 100 SPD, 12.5% duty cycle.
G) dia-PASEF method for 21 min gradients / 60 SPD, 8.3% duty cycle.
H) dia-PASEF method for 44 min gradients / 30 SPD, 4% duty cycle.

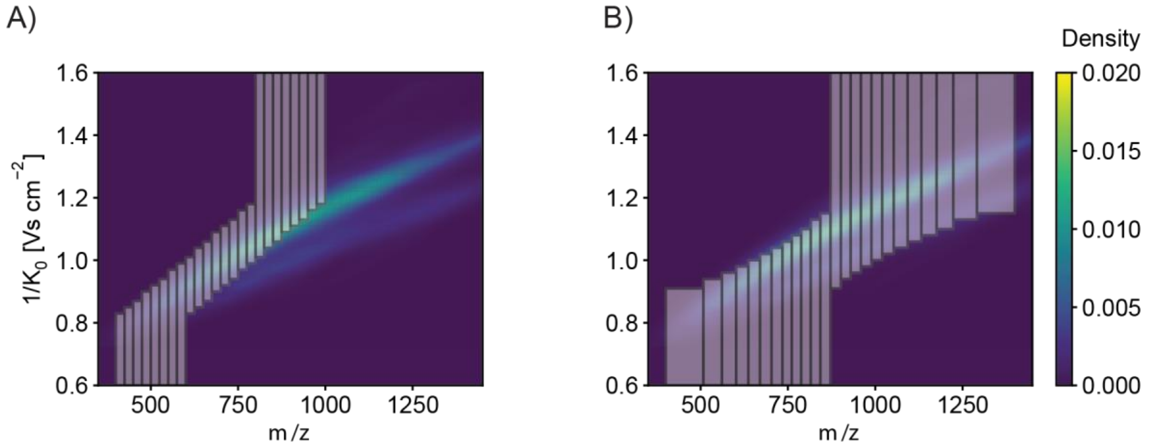


**Supplementary Figure S4: dia-PASEF method plotted on top of kernel density estimation of 'in-depth phospho-library'.**

A) Original dia-PASEF method (6), 12.5% duty cycle.

B) dia-PASEF method for 21 min gradients / 60 SPD and phosphoproteomics, 8.3% duty cycle.

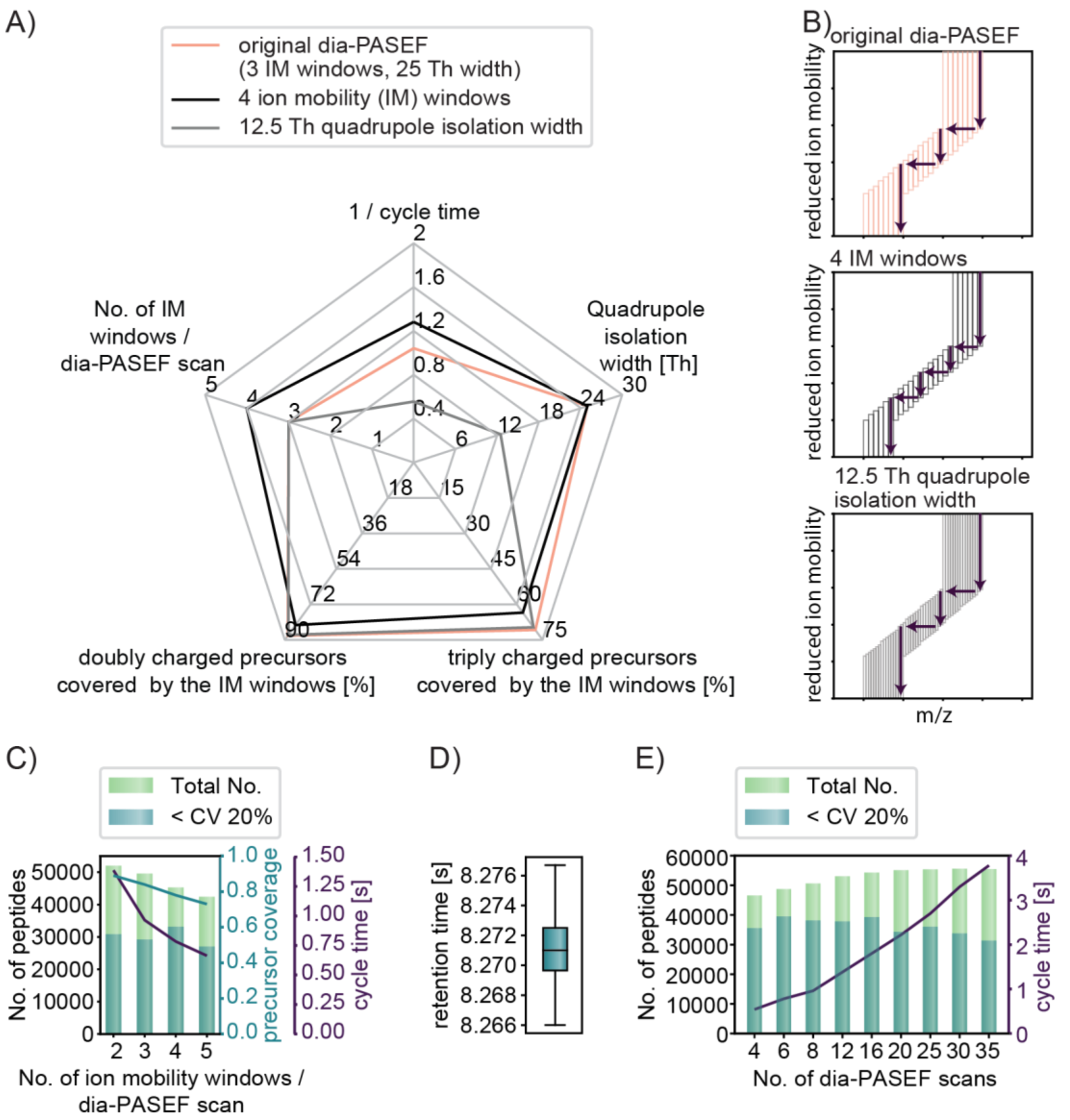


**Supplementary Figure S5: Optimization of the original dia-PASEF method.**

A) Theoretical considerations of the trade-off between identification, quantification, and peptide ion coverage influenced by the number of dia-PASEF scans, number of ion mobility windows per dia-PASEF scan, and quadrupole isolation width.

B) Three exemplary acquisition schemes: original dia-PASEF method (top), original method but with four ion mobility windows per dia-PASEF scan (middle), and original method but with half of the quadrupole isolation widths (bottom).

C) Number of peptides identified and quantified when testing a different number of ion mobility windows per dia-PASEF scans. The method with two ion mobility windows per dia-PASEF scan yields the highest theoretical and empirical peptide coverage.

D) Retention times of precursors with a CV below 20%. Boxplots show the median (center line), $25^{th}$, and 75th percentiles (lower and upper box limits, respectively), and the 1.5× interquartile range (whiskers). n = 44,758 acquired with a method with 12 dia-PASEF scans.

E) Number of peptides identified and quantified when testing a different number of dia-PASEF scans. The number of dia-PASEF scans was individually adjusted to the gradient lengths. Equally good compromises of total identifications and peptide identification with a CV below 20% were obtained at both 12 and 16 dia-PASEF scans. The method with 12 dia-PASEF scans was used in the rest of the study.

The results in C are based on triplicate and in D on quadruplicate injections of 200 ng tryptic HeLa digest with a 60 SPD method. The raw files are searched with the ‘reference proteome library’ as described in Experimental Procedures.

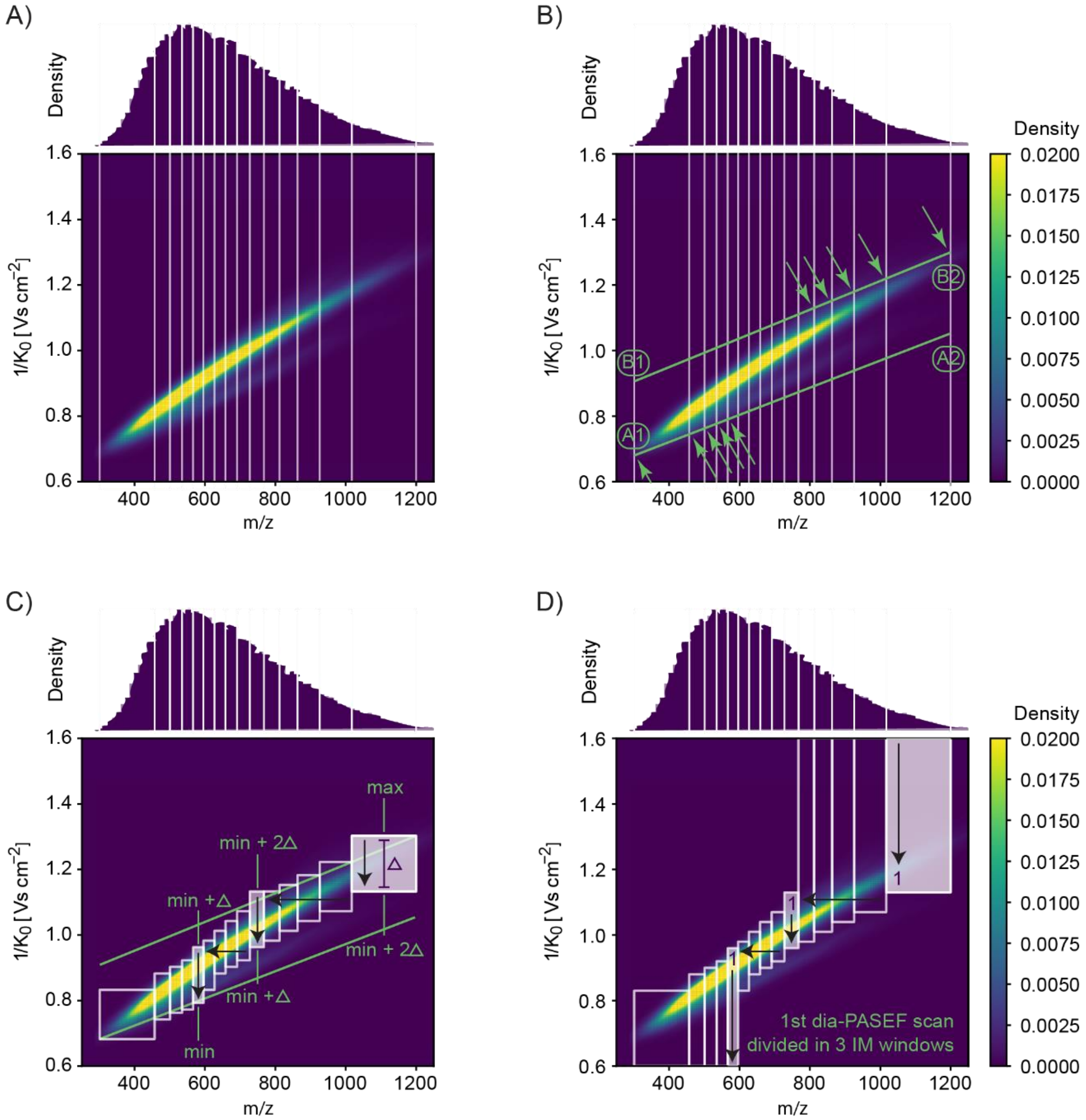


**Supplementary Figure S6: Algorithm of py_diAID in detail.**

A) Binning the *m/z*-IM-plane into isolation windows with an equal number of precursors: n dia-PASEF scans x m IM windows = N bins (e.g., 5 dia-PASEF scans x 3 IM windows lead to 15 bins).

B) The scan area is defined by two diagonals (A1-A2, and B1-B2) The intersections of the bins and the scan area define the position of the acquisition scheme in the *m/z*-IM plane.

C) Calculation of the IM width of each isolation window, Δ = (max - min) / n dia-PASEF scans.

D) Extension of the IM windows to the border of the IM scan range for maximized precursor coverage.

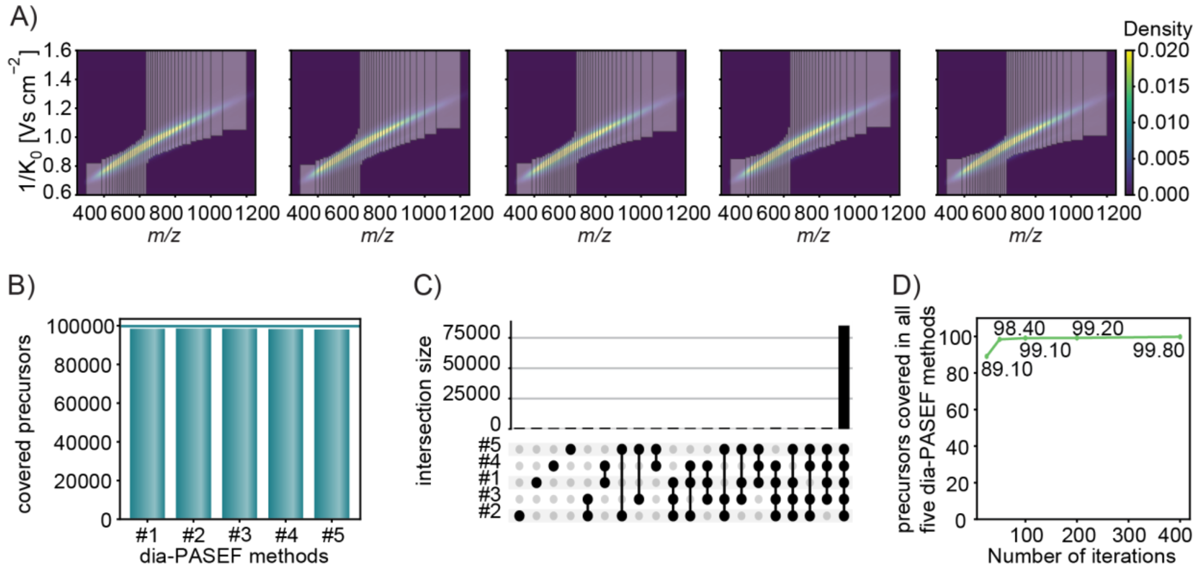


**Supplementary Figure S7: High reproducibility of the py_diAID algorithm.**

A) py_diAID generated five dia-PASEF methods with 200 iterative optimization steps for each method. All five methods are very similar.

B) Similar simulated precursor coverage of each dia-PASEF method. The horizontal line shows the precursor coverage of the library.

C) This upset plot shows precursors that are covered by one or multiple dia-PASEF methods. All five dia-PASEF methods contain 99.2% of the precursors.

D) py_diAID generated five dia-PASEF methods with 25, 50, 100, 200, and 400 iterative optimization steps. The plot depicts the percentage of shared precursors in all five dia-PASEF methods. We found the balance between reproducibility and simulation time at 200 iterations.

All simulations and calculations are based on the ‘reference proteome library’.

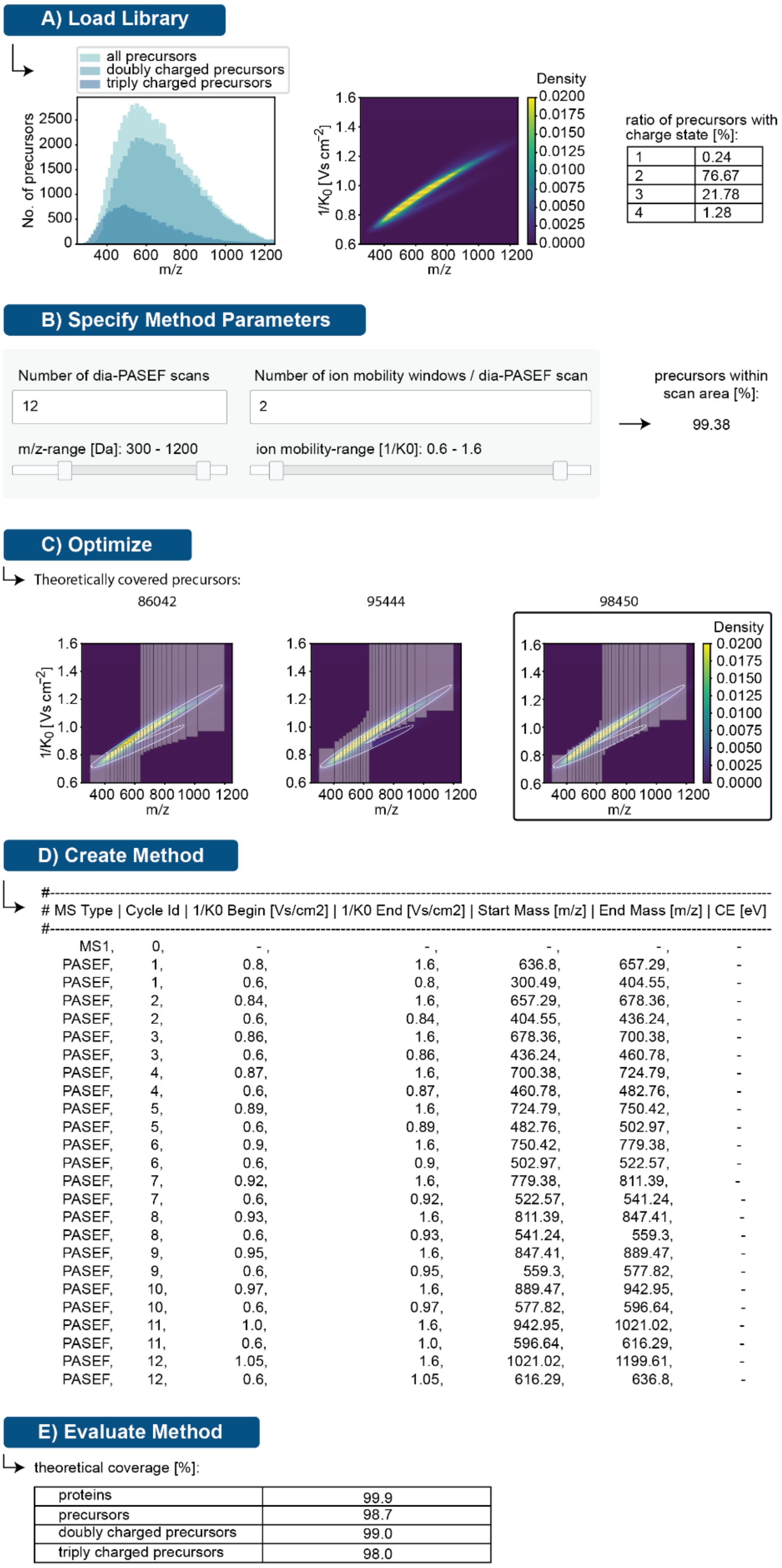


**Supplementary Figure S8: The graphical user interface of py_diAID.**

A) Loading of the spectral library to first evaluate the precursor population of interest by the charge state distribution and the position of the precursor cloud in the *m/z*-IM plane.
B) Definition of the method parameters and selection of the *m/z* and IM limits.
C) Automatic isolation design with Bayesian optimization to find the optimal acquisition scheme.
D) Automatic generation of the parameter file of the optimized dia-PASEF method ready to import into the acquisition software "timsControl" (Bruker Daltonics).
E) Evaluation parameters of the final dia-PASEF method designed by py_diAID.

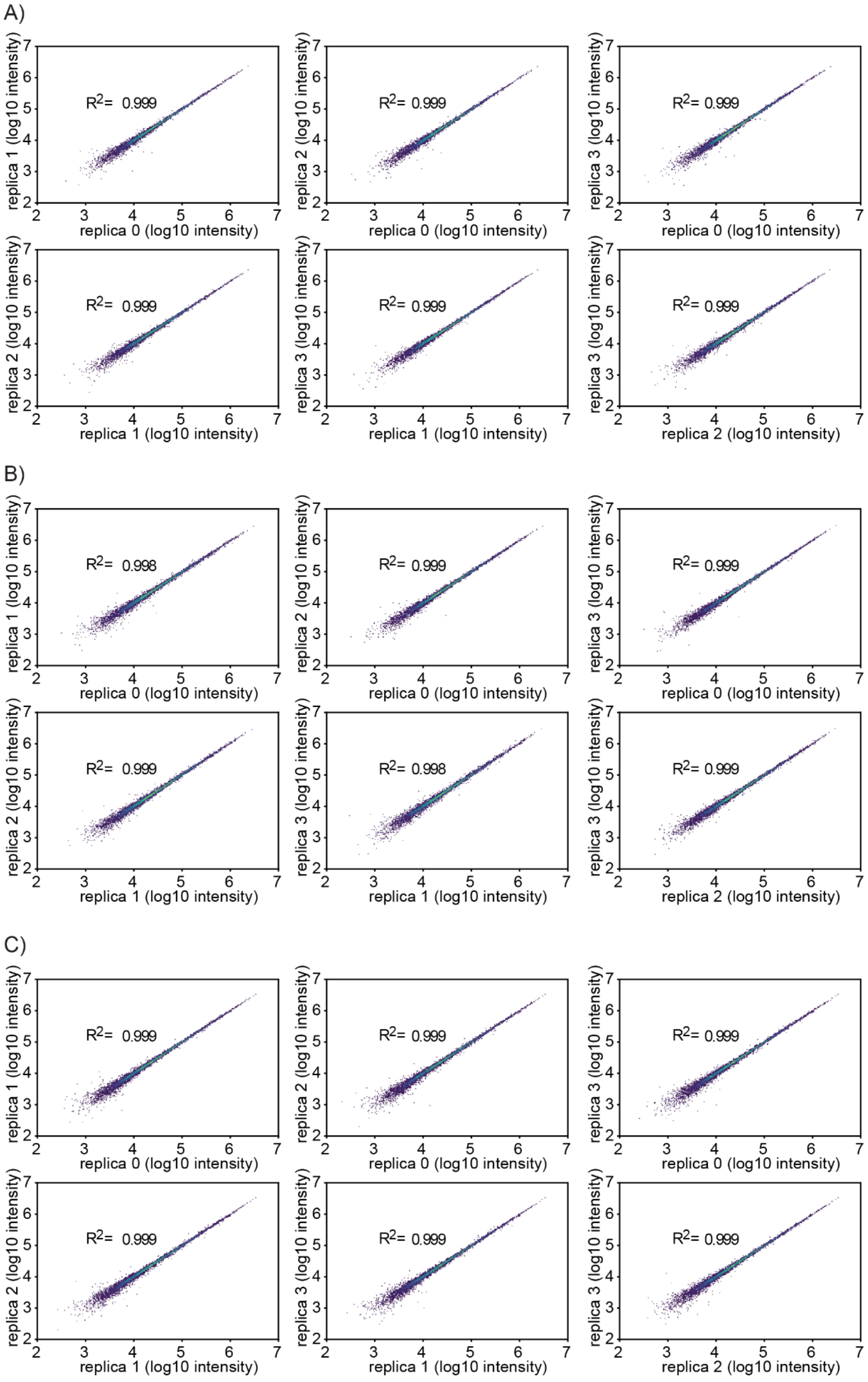


**Supplementary Figure S9: Pearson correlation between the rank ordered quantification values acquired with different gradient lengths. $Log_{10}$-transformed protein intensities on the axis with replica n vs. replica m.**

A) 11-minute gradient.

B) 21-minute gradient.

C) 44-minute gradient.

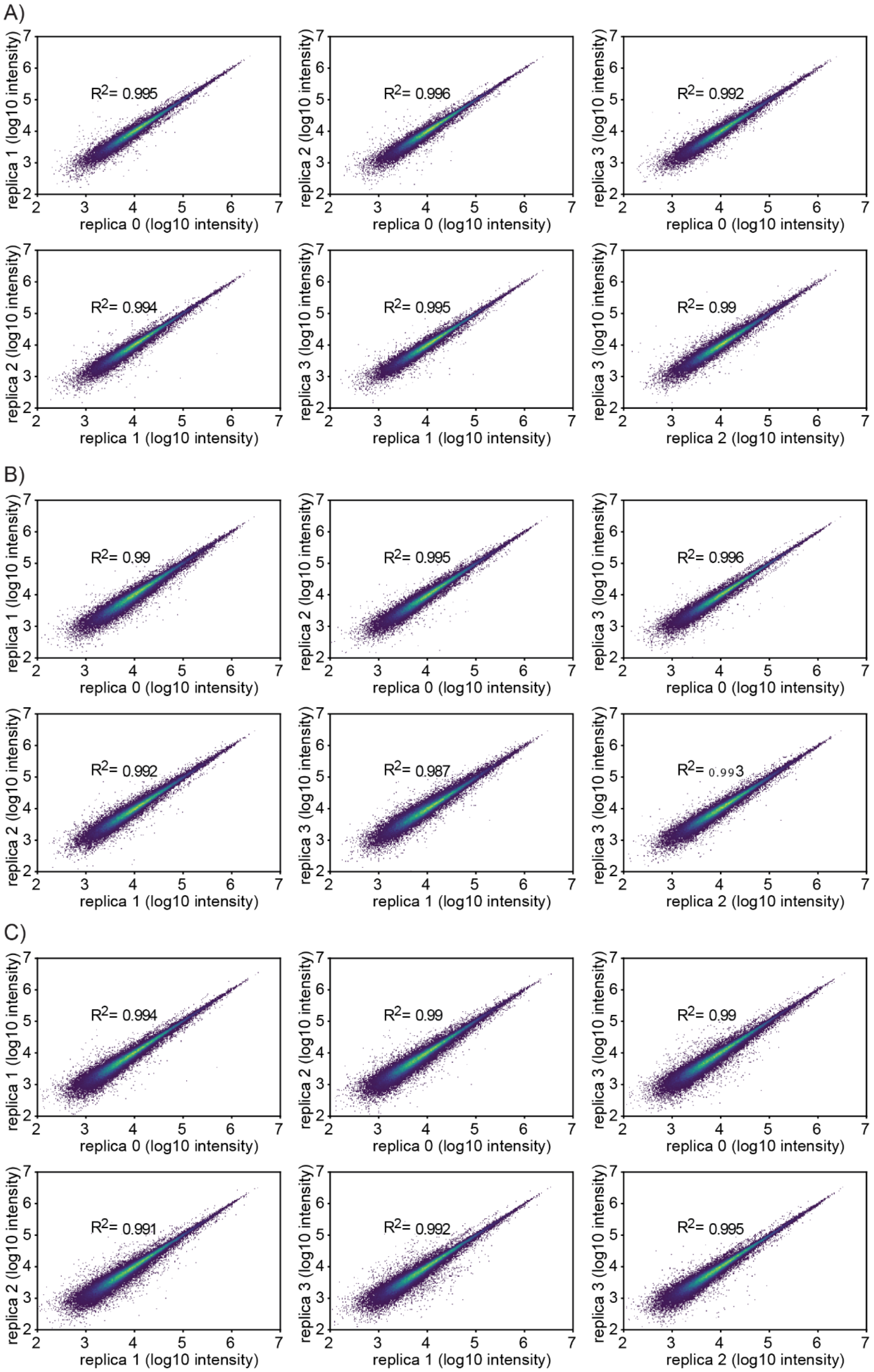


**Supplementary Figure S10: Pearson correlation between the rank ordered quantification values acquired with different gradient lengths. $Log_{10}$-transformed peptide intensities on the axis with replica n vs. replica m.**

A) 11-minute gradient.
B) 21-minute gradient.
C) 44-minute gradient.

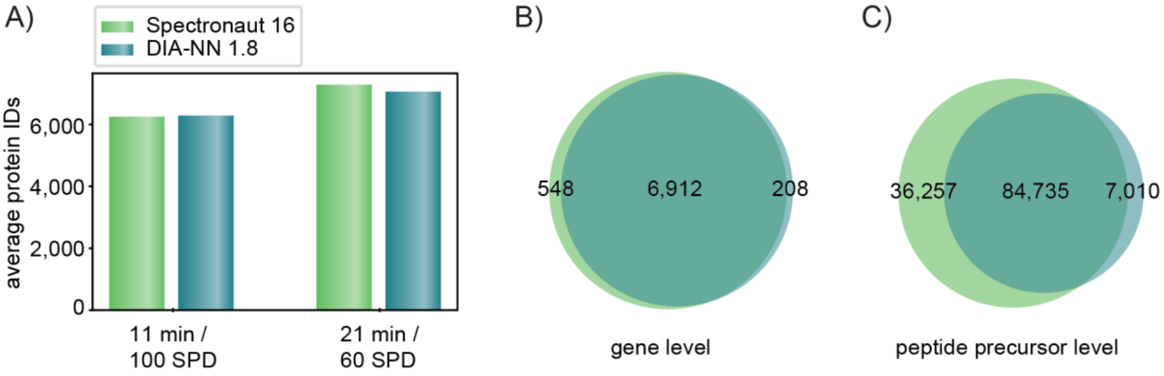


**Supplementary Figure S11: Proteomics analysis with DIA-NN 1.8 vs Spectronaut 16.**

A) Average protein identifications at a throughput of 100 and 60 samples per day (SPD).

B) Overlap between Spectronaut and DIA-NN identifications on gene level within 21 minutes.

C) Overlap between Spectronaut and DIA-NN identifications on peptide precursor level within 21 minutes.

The results are based on quadruplicate injections of 200 ng tryptic HeLa digest. 11-min FragPipe library: 8,553 proteins and 147,525 precursors, 21-min FragPipe library: 8,439 proteins and 148,244 precursors, 11-min Spectronaut library: 8,583 proteins and 139,673 precursors, 21-min Spectronaut library: 8,519 proteins and 139,198 precursors.

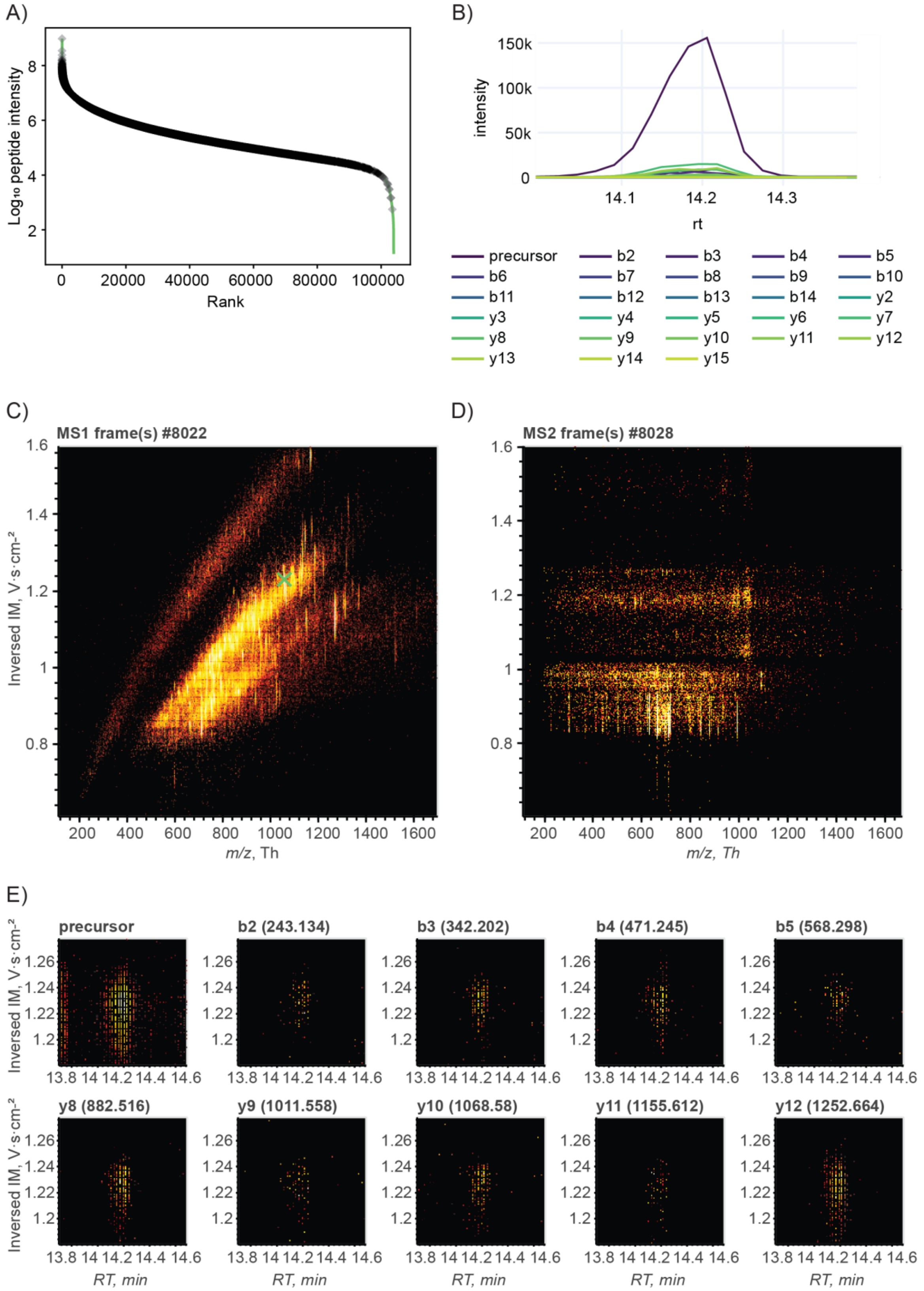


**Supplementary Figure S12: Manual QC of phosphoproteomics data.**

A) Rank plot of the $Log_{10}$-transformed peptide intensities based on the FragPipe phospho-library and DIA-NN single-shot analysis.

B) Elution profile of ELVEPLT[Phospho (STY)]PSGEAPNQALLR.

C) MS1 heatmap including the precursor peak of ELVEPLT[Phospho (STY)]PSGEAPNQALLR.

D) MS2 heatmap including the fragments of ELVEPLT[Phospho (STY)]PSGEAPNQALLR.

E) Precursor and selected fragment peaks of ELVEPLT[Phospho (STY)]PSGEAPNQALLR.

We generated B to E with the automated visualization pipeline AlphaViz (49) that is based on AlphaTims (50).

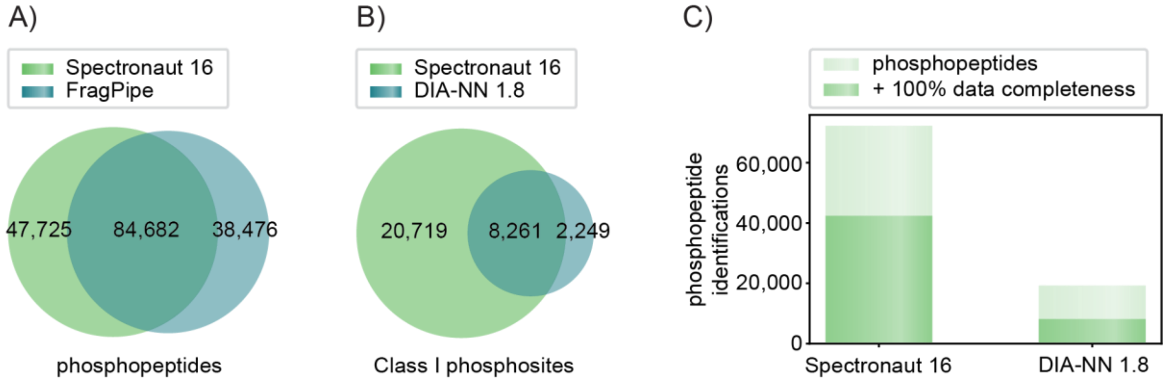


**Supplementary Figure S13: Phosphoproteomics analysis.**

A) Unique and shared phosphopeptides in both phospho-libraries with FragPipe vs Spectronaut 16.

B) Reported class I phosphosites with DIA-NN 1.8 vs Spectronaut 16.

C) Identified phosphopeptides in at least one raw file and with 100% data completeness.

The results of B and C are based on quadruplicates of 100 µg EGF-stimulated and enriched HeLa digest, separated within 21 min.

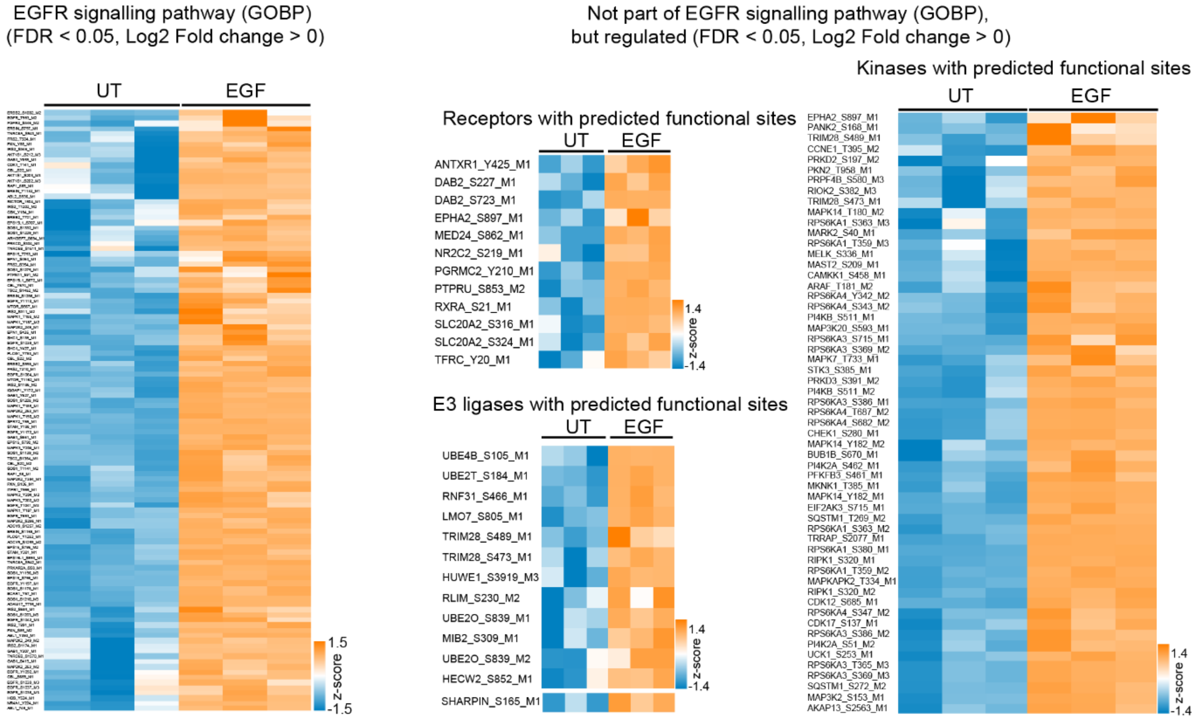


**Supplementary Figure S14: Heatmaps of z-scored intensities of phosphorylation events significantly upregulated upon EGF stimulation.**
